# Host-gut microbiota interactions shape parasite infections in farmed Atlantic salmon

**DOI:** 10.1101/2023.07.20.549827

**Authors:** Jaelle C. Brealey, Miyako Kodama, Jacob A. Rasmussen, Søren B. Hansen, Luisa Santos-Bay, Laurène A. Lecaudey, Martin Hansen, Even Fjære, Lene S. Myrmel, Lise Madsen, Annette Bernhard, Harald Sveier, Karsten Kristiansen, M. Thomas P. Gilbert, Michael D. Martin, Morten T. Limborg

**Author notes:** Address correspondence to: Jaelle C. Brealey, Michael D. Martin, Morten T. Limborg. Morten T. Limborg and Michael D. Martin contributed equally to this work.

## Abstract

Animals and their associated microbiota share long evolutionary histories. Both host genotype and associated microbiota influence phenotypes such as growth and disease resilience. We applied a hologenomic approach to explore the relationship between host and microbiota in shaping lifetime growth and parasitic cestode infection in farmed Atlantic salmon. Genomes, transcriptomes, metabolomes and metagenomes were generated from the guts of 460 harvest-aged salmon, 82% of which were naturally infected with an intestinal cestode. One salmonid-specific *Mycoplasma* dominated the gut microbiota of uninfected salmon. However, the microbiota was perturbed in smaller, parasitised fish, with increased abundance of *Vibrionaceae* and other *Mycoplasma* species previously linked to the cestode microbiota. The cestode-associated *Mycoplasma* carry more virulence-associated genes than the salmonid *Mycoplasma*. Colonisation by one cestode-associated *Mycoplasma* was associated with a region of the salmon genome encoding several long noncoding RNA genes previously associated with host control of intestinal microbiota. Integrating the multiple omic datasets revealed coordinated changes in the salmon gut transcriptome and metabolome that correlated with shifts in the microbiota of smaller, parasitised fish. Our results suggest that cestode infections introduce new microbes and trigger host responses, altering the gut microbiota with increases in potentially pathogenic microbes. Establishment of these microbes is partially shaped by the genetic background of the host. Our study highlights the value of a hologenomic approach for gaining an in-depth understanding of trilateral interactions among host, microbiota and parasite.

## Introduction

There is constant interaction between the vertebrate host and its gut microbiota via the host’s immune system ^1, 2^. The microbiota trains and matures the host immune system ^3^, while the host immune system keeps the microbiota under control to maintain important symbiotic functions, such as providing the host with key nutrients from microbial metabolism of host dietary components ^4^. On one hand, the host genetic background likely affects gut microbiota composition ^5–8^. On the other hand, the gut microbiota has been found to broadly modulate host gene expression by triggering changes in transcription factor binding and chromatin assembly ^9^. Indeed, many studies have demonstrated links between an altered gut microbiota and host disease states, such as infections and inflammatory diseases (reviewed in ^1, 2^). Such disruptions have been referred to as dysbiosis, defined as any change to the resident commensal gut microbiota relative to the community found in healthy individuals ^10^.

Host–microbiota interactions in the gut are particularly relevant to understanding infections by intestinal parasites like helminths (worms), which directly interact with the host immune system and directly or indirectly influence the host gut microbiota ^11^. Gut dysbiosis is frequently associated with helminth infections ^12–14^, and in some systems, the gut microbiota composition directly promotes or inhibits helminth colonisation or reproduction ^15–17^. Furthermore, some parasitic helminths have been shown to carry an internal microbiota ^13, 18–20^, including essential endosymbionts ^21^. Helminths and other parasites may also act as vectors, introducing novel, potentially pathogenic, microbes to the host animal ^22–24^.

The complex interactions among host genotype, host immune system and host microbiota have led to the concept of a “holobiont”, which considers the host organism and its associated microbiota as a single co-functioning scaffold of organisms ^25^. The holobiont has a hologenome consisting of the host genome and the genomes of all its microbiota (the metagenome) ^26^. Thus, it has been argued that the host gut microbiota has to be studied in the context of the host genotype and gene expression landscape to fully understand complex phenotypes, such as lifetime growth or disease susceptibility ^10^. Such hologenomic approaches use multi-omic datasets from both the host (e.g. genomic and transcriptomic) and the microbiota (e.g. metagenomic) to untangle host–microbiota interactions and their associations with phenotypes of interest ^27^.

We explored how host-microbiota interactions shape parasite infections and lifetime growth in a commercial cohort of Atlantic salmon (*Salmo salar*). Production animals often provide excellent systems for hologenomic research because production cohorts are raised under controlled environmental conditions and have well defined phenotypes of interest ^27, 28^. The study salmon population was housed in open seawater pens, allowing natural infection with cestodes (or tapeworms) of the genus *Eubothrium*. This cestode colonises the intestine of the definite salmon host, impeding the salmon’s growth or even causing its death in extreme cases ^29, 30^. We generated and integrated multiple -omic datasets to characterise the gut environment (metagenome, metabolome and host transcriptome) and underlying genomic variability of 460 salmon of differing sizes and cestode infection levels.

## Results

### Summary of datasets

Under our experimental design, we aimed to sample equal numbers of Atlantic salmon across the entire size distribution of a commercial cohort at harvest (gutted weight range: 0.78 – 7.83 kg), resulting in 140 fish classified as ‘small’ (gutted weight ca. < 2.6 kg), 139 ‘large’ fish (gutted weight ca. > 4.2 kg) and 181 ‘medium’ fish (Figure 1A). Salmon were sourced from two sea pens on the same aquaculture farm in southwest Norway fed two different commercial diets (Pen1/Feed1 = 250 individuals and Pen2/Feed2 = 210 individuals). Of these 460 individuals, 375 (81.5%) were parasitised by at least one intestinal cestode. We used a semiquantitative cestode index as a measure of cestode infection load. There was strong evidence that salmon lifetime growth (gutted weight at harvest) decreased as cestode infection levels increased (Figure 1B; p < 0.001, R^2^ = 0.057). Cestode detection was similar between Feed1 and Feed2 (84.4% and 78.1% respectively, p = 0.092). The relative abundance of several important omega-3 and omega-6 fatty acids in muscle tissue were associated with cestode detection, size class and feed type (Supplementary Table 2), including docosahexaenoic acid (DHA), arachidonic acid (ARA), docosapentaenoic acid (DPA) and *alpha*-linolenic acid (ALA).

**Figure 1.**
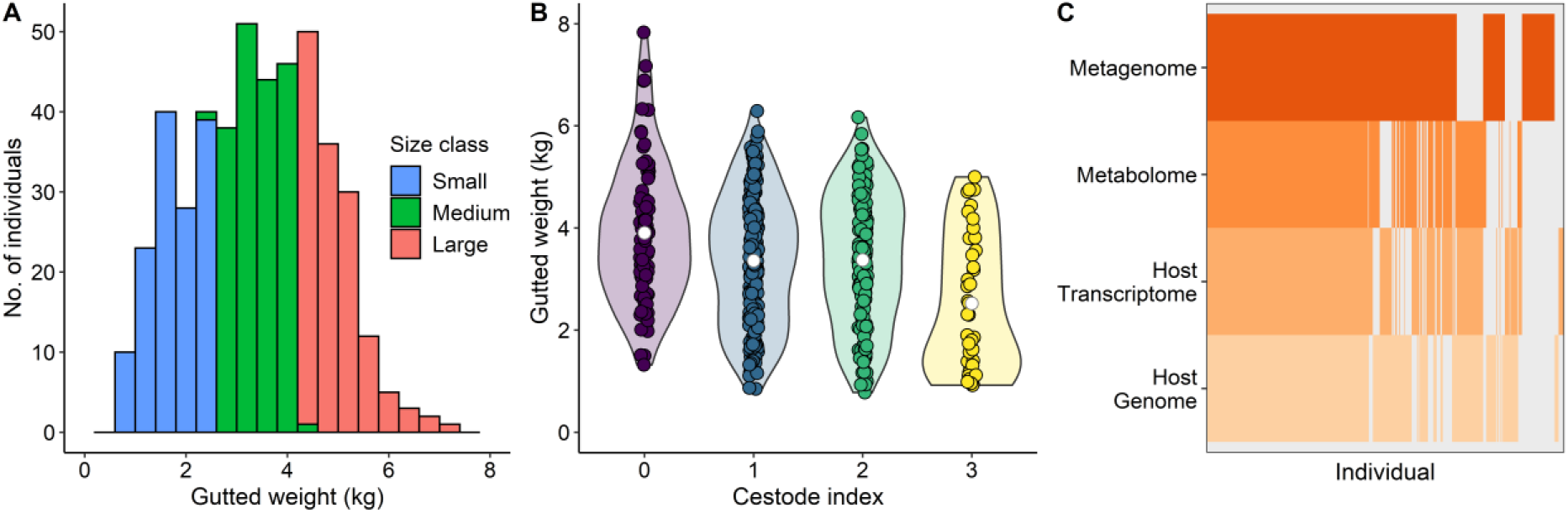
Overview of fish metadata and -omic datasets. **A**) Distribution of lifetime growth of the 460 sampled salmon, measured as gutted weight (kg) at harvest. Individuals were classified into three discrete size classes as indicated by the coloured bars, with the aim to have approximately equal numbers of individuals in each class. **B**) Gutted weight at harvest decreases as cestode infection levels increase. Violin plots show the distribution of salmon weights, white circles indicate the mean. **C**) Gut metagenomes, gut metabolomes, host transcriptomes and host genomes were successfully generated for a subset of individuals. Grey bars indicate that an -omic dataset is missing for that individual. Individuals are ordered by the number of -omic datasets with data available.

After filtering for data quality (see Methods), metagenomes were generated from gut content samples for 392 individuals. Host genomes were resequenced from gill tissue for 361 individuals (coverage average: 4.7X, range: 0.0003–17.3X, 998475 sites after filtering). Transcriptomes from salmon gut epithelial tissue were generated using mRNA sequencing for 343 individuals, resulting in expression data for 19,500 transcripts after processing. Metabolomes, including both salmon and microbial metabolites, were generated from gut content samples for 334 individuals, resulting in an inventory of 969 metabolites, 764 of which could be annotated at the superclass level. A total of 208 individuals were represented in all four -omic datasets (Figure 1C). We therefore aimed to use statistical methods that can accommodate missing data, where possible.

### Cestode infection is associated with an altered gut microbiota

The intestinal microbiota community structure was characterised by low diversity, with only 15 unique bacterial metagenome-assembled genomes (MAGs) recovered from all 392 metagenomic samples (Supplementary Table 1). Most individuals were dominated by MAG01, *Mycoplasma salmoninae* (Figure 2A), a species known to be positively associated with a healthy and undisturbed gut microbiota in salmonids ^19, 31–37^. However, some variation in the microbiota composition was observed in small individuals and those infected with cestodes (Supplementary Figure 1 and Table 3). Alpha diversity was higher in small and/or parasitised fish (Supplementary Figure 2 and Table 4, p < 0.001 and p = 0.014, respectively). The abundance of MAG01 *M. salmoninae* decreased in small fish (Figure 2A, Supplementary Table 4, FDR = 0.0007). Four other *Mycoplasma* species (MAG02-MAG05) were strongly associated with cestode presence (Figure 2B, Supplementary Table 5, FDR < 0.02 for all). Two of these species, MAG02 and MAG03, have been previously associated with the internal cestode microbiota ^19^ (Supplementary Table 1). MAG12 *Carnobacterium maltaromaticum* was also associated with cestode presence (Supplementary Table 5, FDR < 0.05). MAG06 *Photobacterium phosphoreum* was at higher frequency and higher abundance in small fish (Figure 2, Supplementary Table 5, FDR < 0.001). Overall, we observed a shift in the gut microbiota, characterised by decreased abundance of commensal *M. salmoninae* and increased detection of other taxa in small, parasitised salmon compared to larger cestode-free fish.

**Figure 2.**
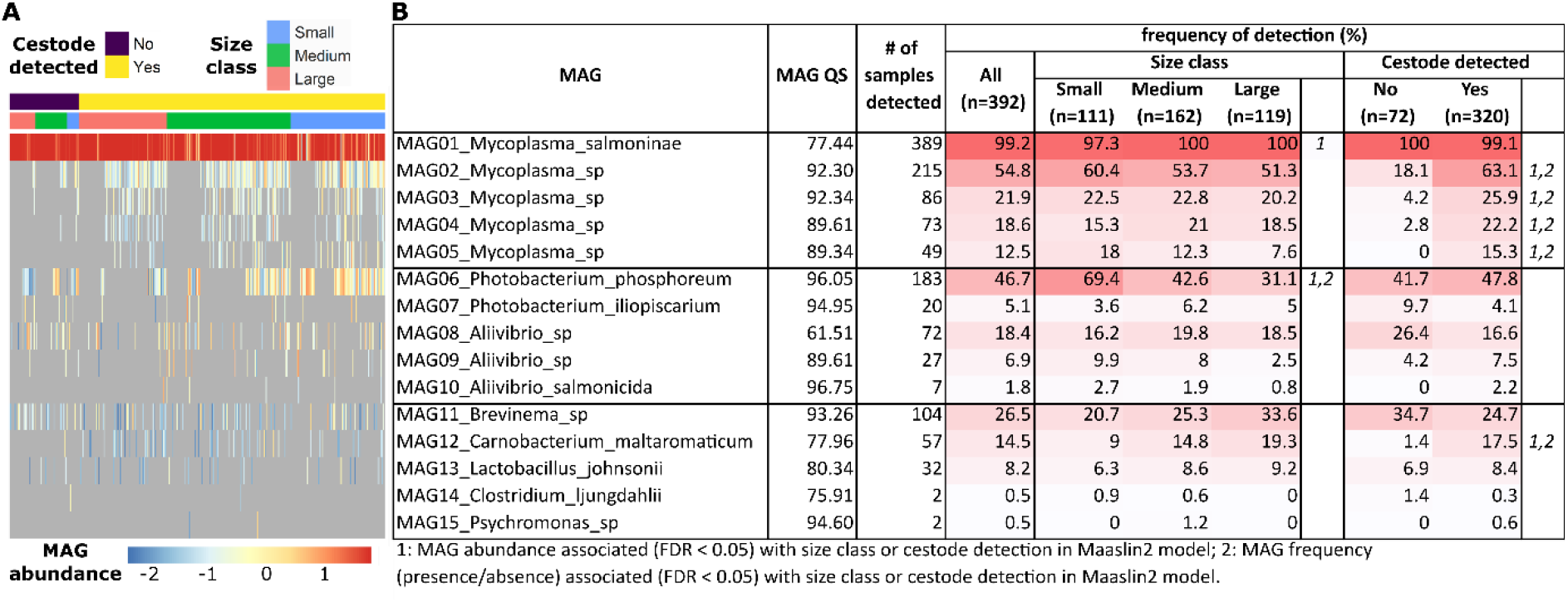
MAG abundance and frequency of detection in the Atlantic salmon gut metagenome. A) Normalised MAG abundances per sample (log-transformed). Samples are ordered and annotated by cestode detection and size class, while MAGs are ordered by taxonomy and frequency of detection. MAGs that were not detected in a sample are shown in grey. B) MAG detection in all samples, separated by size class or cestode detection. MAGs with more frequent detection (% of samples) are coloured in a darker red. MAG QS indicates the quality score of the MAG, based on completeness and redundancy scores (values closer to 100 are better). MAG abundance or detection frequency that was associated (FDR < 0.05) with size class or cestode detection is indicated by *1* or *2*, respectively. Results from the Maaslin2 models with FDR < 0.2 are presented in Supplementary Table 5.

In general, the gut microbiota of the salmon was similar in both feed types (Supplementary Figure 1 and Table 3). However, MAG08 *Aliivibrio* was associated with increased abundance and frequency in Feed2 compared to Feed1 (Supplementary Table 5, FDR < 0.01). Feed type was therefore included as a covariate in all subsequent analyses. We confirmed that the 15 MAGs detected in the salmon gut metagenomes captured the majority of diversity in the gut content at the genus level via 16S metabarcoding of a subset of gut content samples (Supplementary Figure 4). We also compared the gut content samples with paired gut mucosa scrapes and feed pellet samples (Supplementary Figure 3). The most abundant taxa, namely *Mycoplasma*, *Photobacterium*, *Aliivibrio* and *Brevinema*, were detected in both gut content and gut mucosa samples. The only genus detected at high abundance in the feed pellet samples was *Lactobacillus*, which was also detected at low abundance in the gut content samples (Supplementary Figure 3). We therefore conclude that while MAG13 *Lactobacillus johnsonii* may be derived from the feed pellets, the other most abundant MAGs (MAG01–MAG11) are likely true residents of the core salmon gut microbiota.

### Host genomic variation is associated with microbiota composition

We investigated associations between the underlying genomic variability of the salmon host and its gut microbiota composition with a series of genome-wide association studies (GWAS), using both MAG abundance and presence/absence as phenotypes. Cestode detection, size class and feed type were included as covariates. Generally, little host genetic variation was associated with microbiota composition (Supplementary Table 6, Supplementary Figure 5). However, there was one 1.8-Mbp region on chromosome 5 associated with the detection of MAG05 *Mycoplasma* (Figure 3A-B), with two SNPs strongly and 12 SNPs moderately associated with MAG05 detection (p < 5e-8 and p < 1e-5, respectively; Supplementary Table 7). Through linkage disequilibrium analysis we identified an additional 14 SNPs linked to these sites. Of these 28 SNPs, 12 SNPs were located within introns of three coding genes, two SNPs were found in coding regions in two of these genes that resulted in synonymous variants, four SNPs were linked to long non-coding RNAs and 10 SNPs fell in intergenic regions (Figure 3C, Supplementary Table 7). MAG05 was detected more frequently in individuals carrying at least one copy of the minor allele at these sites (Figure 3C). While MAG05 detection was also associated with cestode presence (Figure 2B), cestode presence or absence was not associated with the same host genomic region in the GWAS (Supplementary Figure 5).

**Figure 3.**
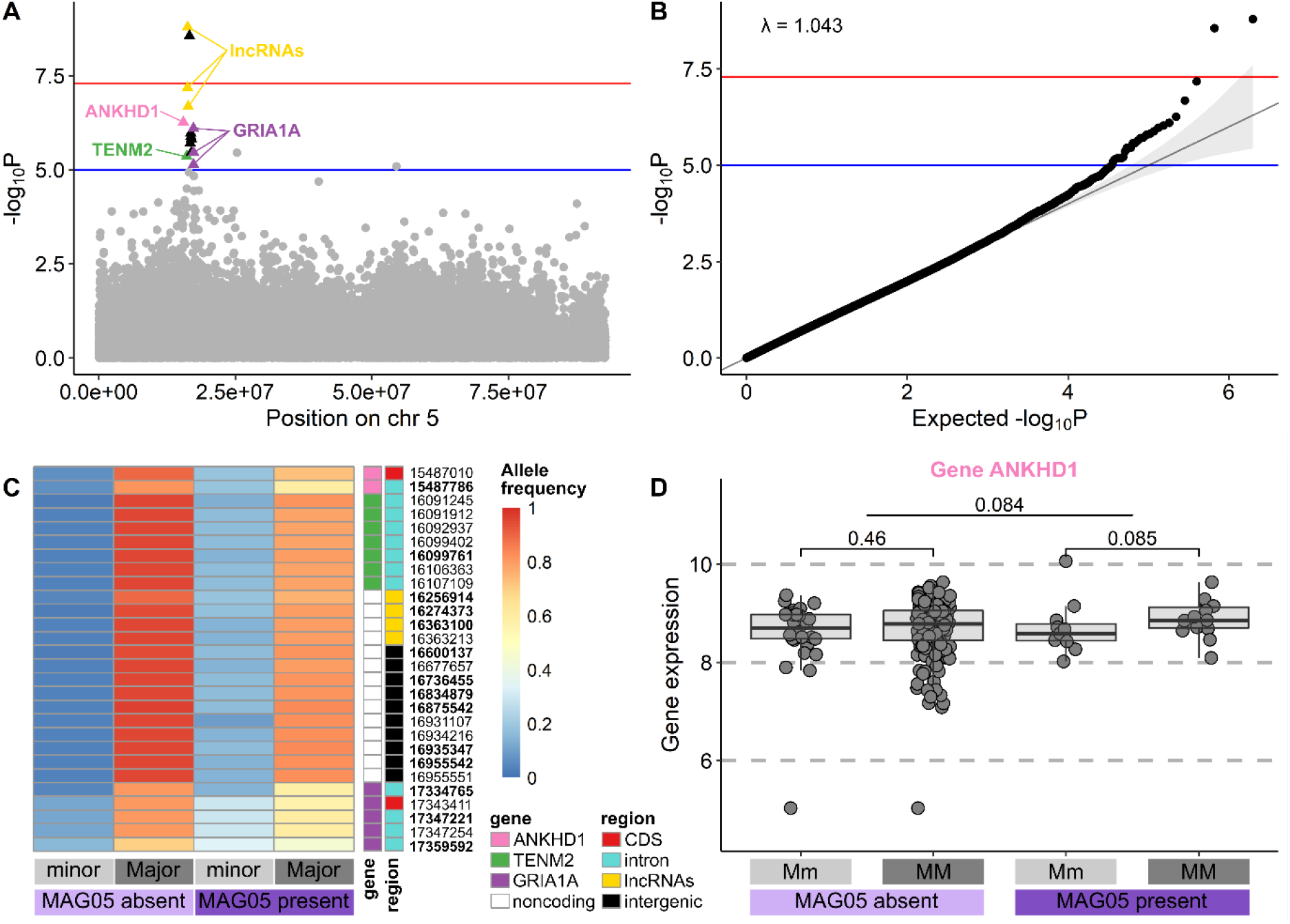
MAG05 Mycoplasma GWAS results. (**A**) SNP peak observed on chromosome 5 for the GWAS association with MAG05 presence/absence. P values have been -log10 transformed so that higher values are more significant. Moderately (p < 1e-5, blue line) and highly (p < 5e-8, red line) associated SNPs on the peak are highlighted in triangles against the background grey circles of chromosome 5. SNPs falling within annotated transcripts in this peak are coloured and annotated by their gene symbol, where relevant (ANKHD1: ankyrin repeat and KH domain-containing protein 1-like; TENM2: teneurin-2-like; GRM1: glutamate receptor 1-like; lncRNAs: long noncoding RNAs). SNPs falling within intergenic regions are shown as black triangles. (**B**) QQ plot and genomic inflation factor (λ) for the GWAS association with MAG05 presence/absence. (**C**) Minor and major allele frequencies for the 28 associated (bold face) and linked (plain face) SNPs. Frequencies have been calculated separately for individuals with MAG05 present or absent. SNPs are annotated by gene symbol (where relevant, same as in A) and genomic region (CDS: coding sequence). (**D**) Gene expression (normalised and scaled counts) of ANKHD1, the only MAG05 associated gene for which comparable transcriptomic data was available. Samples have been divided by MAG05 presence/absence and genotype at SNP 15487786 (Mm: major/minor, MM: major/major; minor/minor individuals were excluded due to low sample numbers). P values are shown for Mm vs MM and MAG05 absent vs present gene expression comparisons using the Wilcoxon rank sum test.

Expression of only one of the three coding genes could be quantified in the mRNA transcriptome dataset (ANKHD1-like). There was weak evidence that ANKHD1-like had increased expression in individuals with the major/major genotype compared to the major/minor genotype (p = 0.084; Figure 3D; only one individual was homozygous for the minor allele, thus this individual was excluded from statistical analysis). This weak association remained in individuals where MAG05 was present, but the trend was not observed in MAG05-negative individuals (Figure 3D). Gene expression analyses of a 1-Mbp region surrounding the chromosome 5 peak in the transcriptomic data did not reveal any differentially expressed genes associated with MAG05 detection or genotype of the 14 SNPs (Supplementary Table 15).

### Multi-omics reveals joint host–microbiota changes in salmon gut environment

We performed multi-omics factor analysis (MOFA) to identify coordinated changes among the salmon gut metabolome, transcriptome and metagenome and correlate these changes with cestode detection and size class variables. MOFA is analogous to PCA, where matrices of - omic data from the same individuals are reduced to a small number of latent factors representing the key contributors of variation across a multi-omic dataset ^38^. Like PCA, factors are ordered by the amount of variance explained (i.e. Factor 1 explains the most variance, etc.). In our MOFA model, we included the 500 most variable features in the metabolome and transcriptome datasets. For the metagenome dataset, we included MAG presence/absence (excluding MAGs detected in < 10 samples), detection of any low-abundant *Mycoplasma* MAG (i.e. MAG02–MAG05), detection of any *Photobacterium* MAG (MAG06–MAG07), detection of any *Aliivibrio* MAG (MAG08–MAG10) and detection of samples with ‘high’ vs ‘low’ abundance of *M. salmoninae* (MAG01; see Methods for details). The model was trained with 15 latent factors. Since feed type explained a small but significant proportion of the variation in the transcriptome and metabolome datasets (ca. 2% each, Supplementary Table 3, Supplementary Figures 6–7), we also repeated the MOFA within each feed type separately. We focus here on consistent results across the three models, i.e., the total dataset, Feed1 or Feed2 only (Supplementary Tables 9-11).

Overall, the MOFA models explained 65–71% of the total variance in the transcriptomic dataset, 48–49% in the metabolomic dataset and 13–22% in the metagenomic dataset (Supplementary Table 8). Generally, each factor in each model explained substantial variation (> 1%) in only one -omic dataset (Supplementary Table 8). However, we identified several factors in each model that explained variation connecting at least two -omic datasets.

The first two factors in each model captured changes generally correlated with size class in either the metabolome or transcriptome (Figure 4, Supplementary Figures 8–9). These changes were usually consistent between feed types. Metabolites associated with large fish included amino acids, peptides, acyl carnitines, fatty acids and fatty acid esters, whereas metabolites associated with small fish included bile acids, hydroxysteroids and derivatives. Gene expression associated with large fish was inconsistent between feed types; however, genes with increased expression in small fish included those with functions in cytoskeleton signalling (e.g. actin and tensin) and immune processes (e.g. the immune modulator thrombospondin-1).

**Figure 4.**
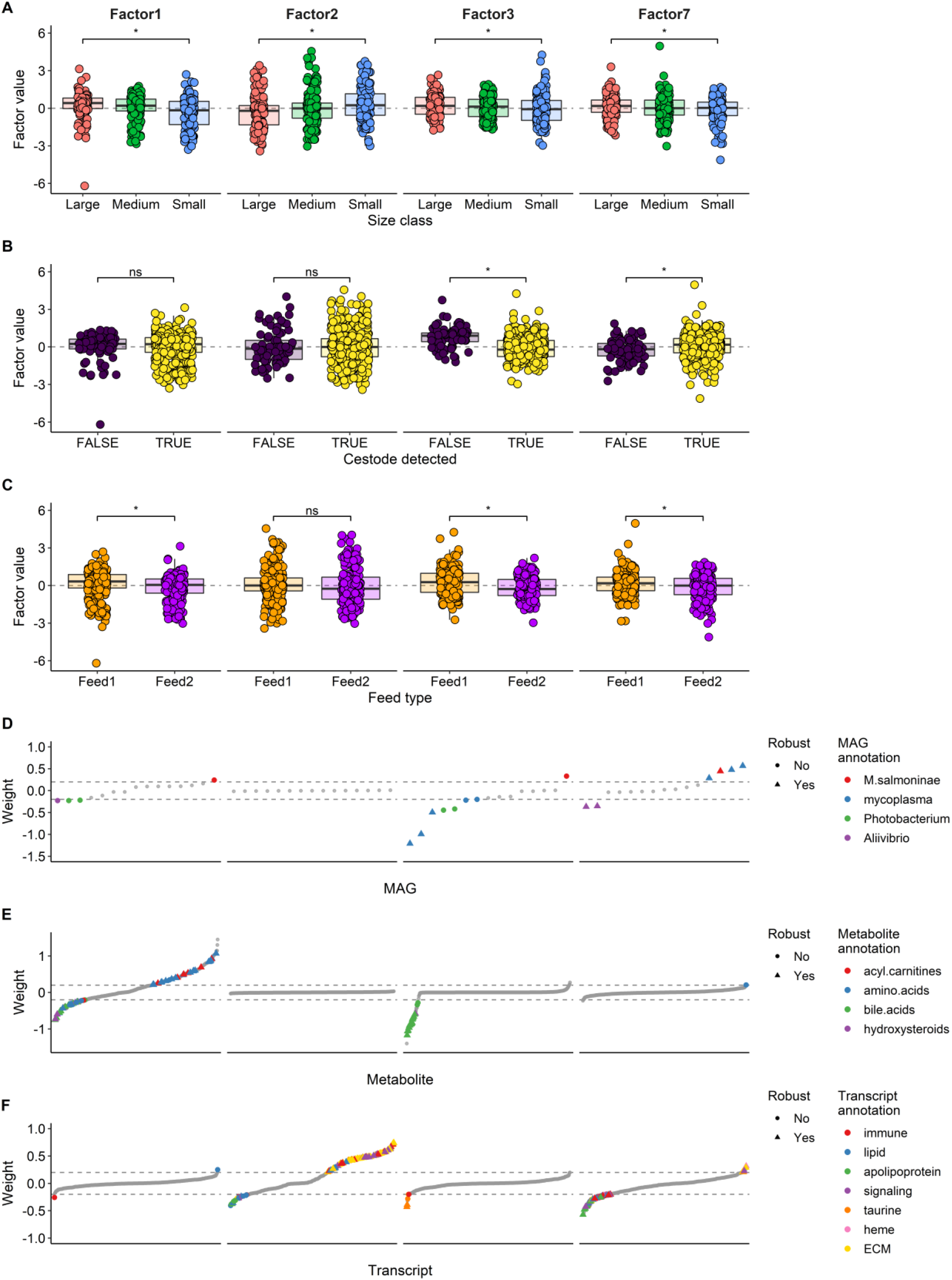
Multi-omics factor analysis (MOFA) results for Factors 1, 2, 3 and 7 in the full model (i.e. both feed types combined). **A**) All four factors were correlated with size class. **B**) Factor 3 and 7 were correlated with cestode detection. **C**) Factors 1, 3 and 7 were correlated with feed type. In A–C, * indicates adjusted p < 0.05, ns indicates adjusted p ≥ 0.05. **D–F**) Feature weights for the metagenome (**D**), metabolome (**E**) and transcriptome (**F**) for Factors 1, 2, 3 and 7. Features are ranked according to their weight. The higher the absolute weight, the more strongly associated a feature is with that factor. A positive weight indicates the feature has higher levels in samples with positive factor values, while a negative weight indicates the opposite. Features with weights > 0.2 or < -0.2 are coloured by MAG species or genus (D) or most frequent functional annotation (E-F), while features with less frequent annotations, or those with weights outside the threshold range (as indicated by the dashed lines) are shown in grey. Features that were consistently found between the two feed type MOFA models to have similar patterns as in the full model (i.e. found in factors with similar fish phenotype correlations and similar metagenome composition patterns) are labelled as ‘robust’.

One factor in each model captured correlated changes in the metagenome and metabolome (Factor 3 in the combined model in Figure 4, Factor 3 in the Feed1 model in Supplementary Figure 8, Factor 4 in the Feed2 model in Supplementary Figure 9). This factor was correlated with cestode detection (Figure 4B). Consistent with previous observations, low-abundant mycoplasma MAGs showed increased detection in parasitised fish, while a high abundance of MAG01 *M. salmoninae* tended to be associated with increased detection in non-parasitised fish (Figure 4D). Several metabolites showing an increased abundance in parasitised fish were annotated as bile acids, alcohols and derivatives (Factor 3 in Figure 4E). While this factor did not substantially contribute to the transcriptome dataset, two genes with core roles in the biosynthesis of taurine, a key component of taurinated bile acids, had consistently higher expression in parasitised fish (Factor 3 in Figure 4F).

We also observed several factors that captured changes generally correlated with size class in the metagenome and transcriptome (Factor 7 in Figure 4). However, these changes were partially dependent on feed type – while MAG01 was associated with large fish in all three models, within Feed1 MAG06 *P. phosphoreum* was associated with small fish (Factors 6-7 in Supplementary Figure 8), while within Feed2 MAG08 *Aliivibrio* tended to be associated with small fish (Factor 5 in Supplementary Figure 9). Generally, antimicrobial peptides, like ladderlectin and cathelicidin, and lipid-binding apolipoproteins had increased expression in small fish while collagen had increased gene expression in large fish (Factors 6-7 in Supplementary Figure 8, Factors 5-6 in Supplementary Figure 9).

### Phylogenetic and functional characterisation of cestode- and size-associated MAGs

To understand how the identified associations have been shaped through interactions between gut microbes and the salmon host, we compared the genomes of the cestode-associated mycoplasma MAGs (MAG02–MAG05) with MAG01 *M. salmoninae* and three related *Mycoplasma* references, using two *Ureaplasma* genomes as outgroups (Supplementary Table 12-13). Average nucleotide identity (ANI) was < 85% between all pairs of genomes (Supplementary Table 12), supporting the conclusion that each *Mycoplasma* MAG forms a separate species. The *Mycoplasma* MAGs clustered into three clades (Figure 5A): MAG01 (*M. salmoninae*) was most similar to *M. iowae* and *M. penetrans*, as previously reported ^19, 31, 32^; MAG03 and MAG05 were more similar to the known fish pathogen *M. mobile* (hereafter referred to as the ‘*M. mobile* clade’); and MAG02 and MAG04 formed a novel, more divergent clade (the ‘MAG02/MAG04 clade’).

**Figure 5.**
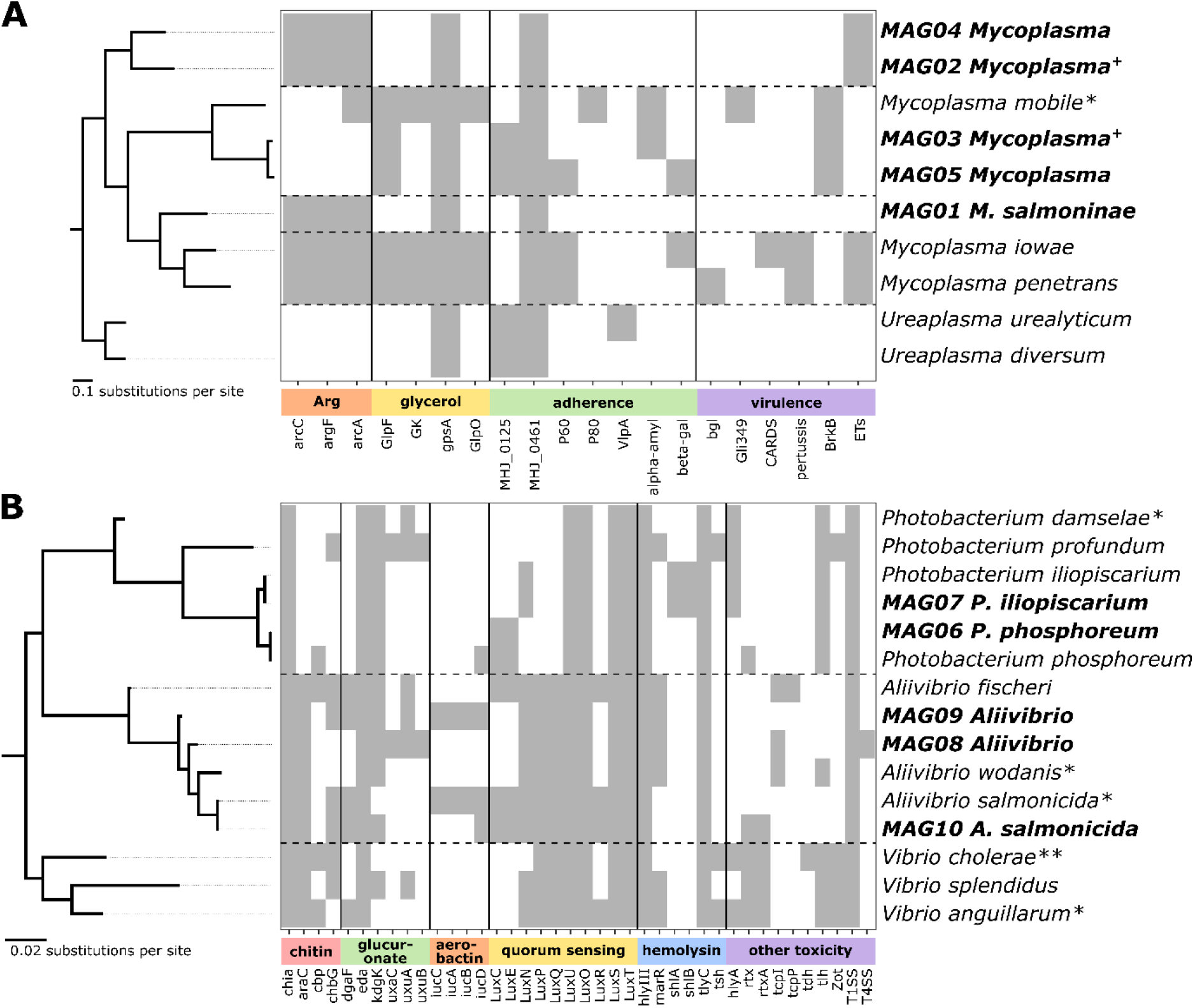
Phylogenetic and functional analysis of *Mycoplasmataceae* (**A**) and *Vibrionaceae* (**B**) MAGs compared to reference genomes. Genomes are ordered by their phylogenetic relationships, from a tree built from single-copy bacterial genes present in all genomes. Trees were rooted using *Ureaplasma* (A) and *Vibrio* (B) reference genomes as outgroups. Branch lengths are measured in substitutions per site. MAGs from this study are highlighted in bold. Heatmap presents the presence (grey) or absence (white) of genes involved in nutrient metabolism, colonisation and/or virulence functions. Genes are grouped by related functions. In A, these are genes in the arginine deiminase pathway (“Arg”), glycerol uptake and metabolism (“glycerol”), adherins that bind to host cell molecules and enzymes which cleave host cell polysaccharides to aid adherence (“adherence”) and more specific virulence factors, including the *M. mobile* gliding mechanism Gli349 and bacterial toxin production (“virulence”). Panel B shows genes of known relevance to the *Vibrionaceae* clade, specifically genes involved in chitin degradation, involved in the colonisation of invertebrates (“chitin”), the D-glucuronate degradation pathway, involved in adaptation to the gut (“glucuronate”), synthesis of aerobactin, a siderophore used to sequester iron (“aerobactin”), community communication via quorum sensing (“quorum sensing”), hemolysin toxin production (“hemolysin”) and other toxin-producing genes (“other toxicity”). A complete list of gene symbols with detailed names and functions is provided in Supplementary Table 14. + after MAG names indicates *Mycoplasma* MAGs previously associated with the body cavity of the cestode, * indicates *Mycoplasmataceae* and *Vibrionaceae* reference species known to be pathogenic in fish and ** indicates *Vibrionaceae* reference species known to be pathogenic in humans.

A number of virulence factors have been characterised in the *Mycoplasma* genus, although much of the focus in the literature is on pathogens of terrestrial animals ^39^. We focused on those factors involved in adaptations for the utilisation of specific nutrients, production of hydrogen peroxide, colonisation and adherence to host cells and toxin production (Figure 5A). As previously reported ^19^, genomes in the *M. mobile* clade lacked the three genes required to generate ATP from the metabolism of arginine, in contrast to *M. salmoninae* and the MAG02/MAG04 clade. All three *M. mobile* clade genomes contained genes required for the uptake and metabolism of glycerol, a potential alternative energy source ^40^. However, only *M. mobile* had glycerol-3-phosphate oxidase (GlpO), which generates cytotoxic hydrogen peroxide through glycerol metabolism. One putative adhesin (MHJ_0461) was conserved across *Mycoplasma* genomes. The *M. mobile* clade genomes also contained additional genes reported to be involved in adherence ^39^. Focusing on other virulence factors, the MAG02/MAG04 clade genomes included genes with sequence similarity to secreted exfoliative toxins, which are produced by *Staphylococcus aureus* to disrupt the epithelial cell layer during skin infections ^41^. In contrast, the *M. mobile* clade genomes included genes similar to the virulence factor BrkB, which is part of an auto-transporter of *Bordetella pertussis* that confers resistance to killing by the classical complement pathway ^42^. Despite the relatedness of the *M. mobile* clade, only the reference *M. mobile* contained the gliding protein Gli349, which provides *M. mobile* with its unique mechanism for gliding motility and is suggested to be important for its pathogenicity ^43^. As previously reported, *M. salmoninae* did not have genes with any similarity to these known *Mycoplasma* virulence factors ^31^.

We performed a similar analysis with the genome of the size-associated MAGs (MAG06 *P. phosphoreum* and MAG08 *Aliivibrio*), compared to the other *Vibrionaceae* MAGs (MAG07 *P. iliopiscarium* and MAG09 *Aliivibrio*) and seven related *Vibrionaceae* references (*P. damselae*, *P. profundum*, *P. iliopiscarium*, *A. fischeri*, *A. wodanis* and *A. salmonicida*), using an additional three *Vibrio* genomes as outgroups (*V. cholerae*, *V. splendidus* and *V. anguillarum*). For all three genera, we aimed to include a range of reference species that included known fish pathogens and commensal or environmental marine species (Figure 5B, Supplementary Table 13).

In the *Photobacterium* genus, MAG06 and MAG07 shared > 98% ANI to the reference *P. phosphoreum* and *P. iliopiscarium* genomes, respectively, supporting their classification as these species (Supplementary Table 12). *P. iliopiscarium* has been previously isolated from the gut microbiota of various Norwegian marine fish species, including Atlantic salmon and has not been associated with disease ^44^. *P. phosphoreum* has been found in association with the skin and gut of healthy salmonids and other fish ^45, 46^. Both species were more distantly related to the free-living seawater species *P. profundum* and the fish skin pathogen *P. damselae* (Figure 5B). As previously reported ^47^, the luminescence genes forming structural parts of the luciferase operon (luxCE) were present in the *P. phosphoreum* genomes, while they were absent from the other *Photobacterium* genomes (Figure 5B). The *P. iliopiscarium* genomes contained more known hemolysins than *P. phosphoreum*, but such toxins were also present in the nonpathogenic *P. profundum*.

In the *Aliivibrio* genus, MAG10 shared 97.6% ANI with the reference genome of the salmonid pathogen *A. salmonicida* (Supplementary Table 12). The *A. salmonicida* genomes grouped with the salmonid skin pathogen *A. wodanis*, rather than the commensal cephalopod symbiont *A. fischeri* ^48^ (Figure 5B). MAG08 and MAG09 shared approximately 85% ANI with each other and with *A. wodanis*, supporting their classification as separate species (Supplementary Table 12). Both MAGs grouped with the putative pathogenic *Aliivibrio* reference species rather than the commensal *A. fischeri* (Figure 5B). Previously, chitin degradation has been associated with *Aliivibrio* commensalism or symbiosis with marine invertebrates, such as copepods or cephalopods ^49^. Consistent with previous reports ^50^, *A. salmonicida* did not have a complete chitin degradation pathway (Figure 5B). MAG08 and MAG09 were also missing key components of this pathway, unlike the cephalopod-associated *A. fischeri*, suggesting that these *Aliivibrio* are not adapted to colonisation of such invertebrates. The luxCE genes required for the bioluminescence associated with *A. fischeri*–cephalopod symbiosis were not observed in MAG08 and MAG09. However, all *Aliivibrio* genomes contained other common quorum-sensing autoinducers and regulators. MAG08 had a complete D-glucuronate degradation pathway, while this pathway was incomplete in other *Aliivibrio*. Gut microbes can use this pathway to utilise host glucuronate as a carbon source ^51^, suggesting a possible adaptation of MAG08 to the salmon gut environment. Some toxins and toxin-related systems, including hemolysin III, tlyC and type I secretion system (T1SS), were conserved across *Aliivibrio* genomes. However, there were no clear patterns between the presence of toxicity genes and known pathogens.

## Discussion

The gut microbiota of the piscivorous Atlantic salmon is characterised by low biomass and low diversity ^32^, and thus is potentially easily disrupted by factors like immune dysregulation, infections, therapeutics and dietary changes ^2^. In our large cohort of farmed salmon, we observed clear differences in the gut metagenomes of large, non-parasitised fish compared to small, parasitised individuals, which correlated with variation in the salmon genome, transcriptome and metabolome. A single *Mycoplasma*, MAG01, dominated large, non-parasitised fish, consistent with previous studies showing high levels of *Mycoplasma* in the gut microbiota of both farmed and wild Atlantic salmon ^19, 31–37, 52^. While small and/or parasitised salmon also had high abundance of MAG01, we observed increased alpha diversity in these individuals, driven by increased frequency of low-abundance *Vibrionaceae* and other *Mycoplasma* MAGs. Our multi-omic approach identified an increased abundance of genes and metabolites associated with the emerging MAGs. Our results therefore suggest that the gut microbiota of small and/or parasitised fish is in a state of dysbiosis, which we discuss in more detail below.

### Cestode-associated Mycoplasma in parasitised fish

Parasitic helminth infections are frequently associated with gut dysbiosis in animals ^13, 14, 53–55^. However, it is often not clear whether prior dysbiosis promotes subsequent parasite colonisation, or whether the parasite triggers dysbiosis, for example by direct interaction with the host microbiota, introduction of new microbes to the host gut or indirect dysregulation of the immune response ^11^. In our study, we identified four *Mycoplasma* MAGs that were strongly associated with cestode detection. Low-abundance *Mycoplasma* MAGs have been identified in the gut metagenomes of wild Norwegian Atlantic salmon colonised by cestodes ^32^. We have also previously identified two of these *Mycoplasma* MAGs closely associated with the body of cestodes from our farmed salmon cohort, specifically MAG02 and MAG03 ^19^ (Supplementary Table 1). Comparing the genomes of the four cestode-associated *Mycoplasma* MAGs with the dominant MAG01, the former contained more genes typically associated with virulence in *Mycoplasma*, particularly the *M. mobile* clade (MAG03 and MAG05). In contrast, MAG01 generally had a reduced genome with fewer genes associated with *Mycoplasma* virulence. Our results suggest that MAG01 is well-adapted to nonpathogenic colonisation of the salmon gut, as previously reported ^31, 52^, whereas we hypothesise that the cestode may introduce novel, potentially more virulent, *Mycoplasma* MAGs into the salmon gut.

In the GWAS, we identified one strong association between the detection of cestode-associated *Mycoplasma* MAG05 and a 1.8-Mbp region of the salmon genome that encoded several long noncoding RNAs (lncRNAs) and three annotated genes. While the direct effects of these variants are currently unclear, lncRNA expression profiles have been previously associated with specific gut microbiota compositions in mice ^56^, possibly through their involvement in inflammation and other immune responses ^57–59^. Thus, different lncRNA genetic variants may have altered expression profiles, suggesting a possible mechanism for host control of the microbiota. Given that MAG05 was strongly associated with cestode detection, we hypothesise that during alteration of the gut microbiota by cestode infection, the ability of MAG05 to establish in the salmon gut may be at least partially determined by the genetic background of the host. Our transcriptome dataset only included mRNA transcripts, thus we were unable to investigate lncRNA expression patterns. However, we did observe one gene in the region of interest on chromosome 5, ANKHD1-like, with weakly altered expression when MAG05 was present, depending on host genotype. Future studies using e.g. whole transcriptome long-read RNA sequencing, would do well to further investigate these lncRNA loci in order to shed more light on links between noncoding RNA expression patterns and microbiota composition. While the mechanisms may be unclear, our results suggest biological control of the microbiota by the host, explained here by host genotype. Similar associations between SNPs in specific host genes and abundance of specific gut microbes have been observed in recent, large-scale human studies ^5, 6^. Cestode alternation of the gut microbiota has also been shown to depend on host genotype in sticklebacks ^60^, further supporting our results.

The MOFA revealed a clear response in the gut environment during cestode infection, with increased frequency of cestode-associated *Mycoplasma* MAGs, increased abundance of bile acid metabolites and altered expression of two genes in taurine metabolism pathways. Bile acids are involved in a range of important functions and processes in fish, including the digestion of lipids, vitamins and carotenoids, cholesterol regulation, inflammation regulation, intestinal barrier function and antimicrobial activities ^61^. Primary bile acids are produced from cholesterol in the liver, where they are conjugated with predominantly taurine to form bile salts, before transport and excretion into the intestine ^61^. Some members of the gut microbiota can deconjugate and further modify these bile salts into secondary bile acids, which can have additional effects on both the host and specific gut microbes ^62, 63^. Cestodes may use host-derived bile acids or salts as a nutrient source ^64^; however, bile acids may also limit core cestode metabolic pathways ^65^. In Atlantic salmon, lower levels of body cholesterol and bile acids have been associated with increased intestinal inflammation, suggested to be linked to changes in dietary taurine and cholesterol ^66^. As our results were independent of feed type, we suggest that bile acids play an important role mediating host–microbiota–parasite interactions in our system regardless of diet.

### Size-associated Vibrionaceae in small fish

Variation in growth rates in salmon raised under identical conditions can be due to multiple biotic factors, including differences in growth hormone production, appetite, feed conversion efficiency, response to stress, behaviour and infectious disease burden ^29, 67–74^. Selective breeding programs have increased domesticated salmon growth rates ^75^; however, since growth traits are polygenic, understanding the genetic basis of growth in salmon remains a complicated challenge ^76, 77^. Links between the gut microbiota and salmon growth have mostly been investigated in relation to the effect of different diet compositions ^52, 78–81^. Thus, our study is one of the first to investigate how salmon genetic variation and gut microbiota composition may affect growth in an integrated manner, independent of diet.

While we found no strong associations between size-associated MAGs and host genetic variability in the GWAS, we observed a correlated set of size-associated changes in the salmon metagenome, transcriptome and metabolome, after controlling for parasite presence. Some of the most important size-associated microbes were *Vibrionaceae* MAGs, including species of *Photobacterium* and *Aliivibrio*, which were more often detected in small fish. Both *P. phosphoreum* (MAG06) and *P. iliopiscarium* (MAG07) are known to colonise salmonid guts in the absence of disease ^44–46^, and the genomes of these MAGs were not enriched for known virulence factors. In contrast, an increased abundance of *Aliivibrio* species in the gut microbiota has been previously found during bacterial infections in farmed Atlantic salmon ^33, 82^, and *A. salmonicida* (MAG10) is a known salmonid pathogen ^83^. However, we observed that the most abundant *Aliivibrio* (MAG08) carried no additional virulence factors, although its glucuronate degradation pathway suggests adaptation to survive in the salmon gut. Thus, from our results it is unclear whether these *Vibrionaceae* species are directly pathogenic or simply opportunistic colonisers during dysbiosis.

Our multi-omic dataset enabled us to generate new connections between host size and immune response to particular microbes. In the MOFA, we identified a set of transcripts that were associated with size class and the detection of *Vibrionaceae* MAGs, although the genus/species of MAG varied with feed type. Many of the genes with increased expression in small fish colonised by *Vibrionaceae* encode proteins in immune pathways, including ladderlectin, cathelicidin and apolipoproteins. Ladderlectin is an antimicrobial lectin in teleost fish that has been shown to bind LPS from gram-negative bacteria, like *Vibrionaceae* species, in rainbow trout ^84, 85^. The Atlantic salmon ladderlectin genes share significant homology with rainbow trout, suggesting a similar function ^85^. Cathelicidins are antimicrobial peptides that have been shown to have an important role in the response to the gram-negative salmonid pathogen *Yersinia ruckeri* ^86^. Apolipoproteins bind lipids and are therefore involved in lipid transportation ^87^; however, they are also involved in innate immunity in teleosts against both gram-negative and gram-positive bacteria, as well as parasites ^88–91^. Furthermore, an increased abundance of *Photobacterium* has been associated with decreased expression of gut barrier function genes in farmed Atlantic salmon ^92^. Our results are therefore consistent with an innate immune response in small fish, triggered by gram-negative bacteria.

MOFA analysis also revealed that the majority of variation in the salmon gut transcriptome and metabolome (> 20% in each) was explained by factors strongly correlated with size class, independent of microbiota composition. Metabolites consistently associated with large fish included acyl carnitines and fatty acids, which have important roles in energy production and storage, among other functions ^93, 94^. We identified a set of genes whose expression was associated only with small fish, with roles in cell division and cytoskeleton and extracellular matrix signalling, such as actin, tensin and myosin. Changes in myosin gene expression have particularly been linked to changes in growth rate in farmed Atlantic salmon ^95^. The gene set also included thrombospondins, immune genes that interact with the extracellular matrix and inhibit angiogenesis, an important process in growth and development ^96^. Our results demonstrate that different growth-related metabolic and development processes occur in fish with different lifetime growth phenotypes.

### Conclusion

Our hologenomic approach to study gut health in farmed Atlantic salmon revealed a clear association between cestode infection, lifetime growth and gut dysbiosis. This dysbiosis was associated with altered expression patterns in the salmon transcriptome and metabolome, including changes in immune, taurine synthesis and bile acid pathways. We suggest two nonexclusive hypotheses to explain these findings. Firstly, the cestode may colonise the salmon gut, altering the gut microbiota and resulting in an altered host response, possibly with changes to host growth phenotypes. Secondly, small fish may be already stressed with a dysregulated gut environment, resulting in lowered host control of the intestinal microbiota, allowing parasites and/or putative opportunistic pathogens to establish or even outgrow commensals in the salmon gut. Furthermore, our results suggest that the genetic background of the salmon may in part determine how the gut microbiota changes during dysbiosis. These results highlight the value of the hologenomic approach for understanding how the response of an intestinal microbiota community to parasite infection may depend on the genetic background of the host organism.

## Methods

### Experimental design and sampling

Samples used for this study were obtained as part of the HoloFish project (Norwegian Seafood Research Fund, project no. 901436). This cohort has been described previously ^19^. Briefly, we sampled 460 ready-to-harvest Atlantic salmon from a commercial production site close to Bergen, Norway, owned by Lerøy Seafood Group during April 2018. Samples were obtained from two groups reared in separate sea pens and fed two different standard commercial diets (‘Feed1’ and ‘Feed2’). These diets have been anonymised but were manufactured respectively by BioMar and EWOS in 2018. For Feed1, salmon were fed BioMar’s “Energy X” from sea transfer until they were 1 kg in weight, “Power 1000” from 1 kg to 2.5 kg and “Power 2500” from 2.5 kg to slaughter. For Feed2, salmon were fed “EWOS Rapid” from sea transfer until they were 1 kg in weight and “EWOS Extra” from 1 kg to slaughter. A number of relevant metadata variables describing various phenotypes of each fish were recorded (Supplementary Table 16), including gutted weight (kg) and intestinal cestode infection status using a cestode index from 0 to 3 where: 0 = no observed cestode; 1 = one to three visible cestodes; 2 = more than three cestodes but digesta volume was higher than cestode volume in the gastrointestinal tract; and 3 = excessive numbers of cestodes dominating the gastrointestinal volume that were likely impeding the passage of feed along the gastrointestinal tract. Fish were further binned into size classes (small, medium and large) based on gutted weight and cestode detection classes (present, absent) using the above-mentioned cestode index. We used a linear model to test for significant differences (p < 0.05) between gutted weight and cestode index, including feed type as a covariate. We used Fisher’s Exact Test to compare frequency of cestode detection between the two feed types.

Six biological samples were taken from each fish, including muscle tissue for fatty acid profiling, gill tissue for host genomics, gut epithelia for host transcriptomics, gut epithelial cell scrapes for 16S metabarcoding and two gut content samples for metagenomics and metabolomics. We also sampled 10 feed pellets for each feed type and characterised their microbiota using 16S metabarcoding.

### Host genomics

Between 10 and 20 mg of salmon gill tissue were used for DNA extraction, using Qiagen DNeasy® blood & Tissue Kit 96, following manufacturer’s recommendations. Genomic DNA was fragmented using a Covaris LE220+, aiming for 350 bp. For library preparation, 100 ng of DNA per sample were used in a single-tube library preparation method ^97^. Libraries were quantified with qPCR to determine the required number of cycles needed for indexing PCR. Purified libraries were indexed and amplified using customised index primers for MGISeq-2000 ^98^. Indexed libraries were purified using magnetic SPRI-beads (Deangelis, Wang, and Hawkins 1995). Prepared libraries were shipped on dry-ice to BGI-Shenzhen for sequencing on MGIseq-2000, using PE 150 bp chemistry.

Adapter and quality trimming was performed using AdapterRemoval v2.2.4 ^99^, trimming consecutive bases with quality scores < 30 and removing reads with > 5 Ns or < 100 bp after trimming. Duplicate reads were removed with SeqKit rmdup ^100^. Reads were aligned to the Atlantic salmon reference genome (GCA_905237065.2) with bwa mem v0.7.16 ^101^. Paired reads mapping to the reference were retained with samtools v1.9 ^102^. Multiple BAM files from the same individual (including host-mapping reads from the metagenomic sequencing) were merged with samtools merge and mapping duplicates removed with Picard MarkDuplicates v2.25.0 (https://broadinstitute.github.io/picard/). To identify callable regions of the genome, 10 samples were randomly selected and combined into a single BAM file with samtools merge. Read groups were reformatted with Picard v2.20.2. Coverage statistics were evaluated with samtools depth. Based on these values, callable loci were identified using GATK CallableLoci v3.7 ^103^, using a minimum depth of 26 and a maximum depth of 158. Genotype likelihoods were then estimated for all samples with the SAMtools model in ANGSD v0.933 ^104^, using the identified callable loci regions. From these, genotype probabilities were imputed with Beagle v3.3.2 ^105^ and converted to genotype dosages. Linkage disequilibrium pruning was performed with PLINK v2 (http://pngu.mgh.harvard.edu/purcell/plink/), removing all correlated SNPs with R2 > 0.5.

### Host gut transcriptomics

Host transcriptomes were extracted using 20 mg of gut epithelia with Quick-RNA™ Miniprep Kit (Zymo Research), following manufacturer’s recommendations. Prior to extraction, samples were washed with PBS buffer to remove residual RNAlater buffer. Subsequently, samples were shipped on dry-ice to Novogene for polyA enrichment and mRNA library preparation and sequencing. Briefly, messenger RNA was purified from total RNA using poly-T oligo-attached magnetic beads. Purified RNA was fragmented via sonication. The first strand of cDNA was synthesized using random hexamer primers, followed by a second cDNA synthesis. On this cDNA a library was then prepared via A-tailing, adapter ligation, PCR amplification, size selection and purification. The resulting purified library was quantified with a Qubit Fluorometer and real-time PCR, and an Agilent BioAnalyzer 2100 was used to determine the fragment size distribution. The quantified libraries were pooled and 150 bp paired-end reads were sequenced on an Illumina NovaSeq 6000 instrument.

Sequence quality of raw RNA-Seq data was assessed using FastQC v0.11.3 (http://www.bioinformatics.babraham.ac.uk/projects/fastqc/). Quality trimming was performed using AdapterRemoval v0.20.4 ^99^ to remove base pairs with a Phred score < 20 and trimming of poly-A tails > 8 bp. Sequences shorter than 55 bp and all unpaired reads were excluded from subsequent analyses. The quality of trimmed sequences was checked again using FastQC v0.11.3. Reads were then aligned to the Atlantic salmon reference genome (GCA_905237065.2) using STAR aligner ^106^ and default parameters. As RNA degradation was present in most samples, aligned reads were used to generate a gene-specific count matrix across samples using DegNorm ^107^. DegNorm adjusts the read counts for transcript degradation heterogeneity while controlling for the sequencing depth. Likely due to the RNA degradation, many samples had low mapping statistics to the reference genome, thus we excluded all samples with < 50% of reads aligned to the reference. To reduce noise in the dataset, we also only included transcripts with at least 10 counts in at least 50% of the samples included in downstream analyses.

We used the R package DESeq2 ^108^ to estimate library size factors, using ‘shorth’ as the locator function from the genefilter R package, which is the shortest interval that covers half of the adjusted count values in each sample. For unsupervised analyses (ordination, MOFA), we used the DESeq2 function vst to normalise adjusted counts by the above size factors, estimate gene dispersions and apply the Variance Stabilising Transformation (VST) to make the dataset approximately homoskedastic and bring the values into log2 space. This transformed dataset was then used as input for the R function prcomp to generate a PCA. PERMANOVA was performed on Euclidean distances of the transformed dataset using the adonis2 function in the R package vegan v2.6-2 (https://github.com/vegandevs/vegan), testing for associations with feed type and size class (Supplementary Table 3). The transformed dataset was also used for feature selection in MOFA (see below).

For differential expression analysis, the DESeq function was applied to the adjusted count data, with negative binomial generalised linear models and Wald significance tests. DESeq automatically incorporates size factors and gene dispersions into the models. Significantly differentially expressed genes (adjusted p < 0.05) between various fish phenotype variables (e.g. size class, feed type, host genotype – detailed below) were extracted from the results.

### Gut metagenomics

DNA was extracted from approximately 100 mg of each gut content sample using the ZymoBIOMICS™ 96 MagBead DNA Kit (D4308) (Zymo Research), following manufacturer’s recommendations. Before extraction, samples were resuspended in 1X Shield™ (Zymo Research). To minimise batch effects all samples were randomised before any laboratory processing. We also included seven negative controls containing only Shield™ (Zymo Research) in the extractions, in order to identify putative lab reagent contaminants in downstream microbial community analyses. Libraries were initially prepared for BGI sequencing, following the same single-tube library preparation method ^97^ as for the host genome. Prepared libraries were shipped on dry-ice to BGI-Shenzhen for sequencing on MGIseq-2000, using PE 150 bp chemistry. To expand the sample number and sequencing depth for the metagenome, aliquots of the DNA extracts described above were also shipped on dry-ice to Novogene for Illumina library preparation and PE 150 bp sequencing.

Adapter and quality trimming was performed using AdapterRemoval v2.2.4 ^99^, trimming consecutive bases with quality scores < 30 and removing reads with > 5 Ns or < 100 bp after trimming. Duplicate reads were removed with SeqKit rmdup ^100^. Reads were aligned to the Atlantic salmon reference genome (GCA_905237065.2) with bwa mem v0.7.16 ^101^. Unmapped paired-end reads were extracted from the BAM files with samtools v1.9 ^102^. Data from both BGI and Illumina were combined to generate the MAG catalogue. Both co-assembly and single-assembly approaches were used. For co-assembly, samples were divided by feed type. Samples within each group were co-assembled with Megahit v1.2.9 ^109^, using a minimum contig length of 1000 bp in the mode meta-sensitive. Reads from all samples and feed types were aligned back to each co-assembly with bwa v0.7.17. Binning was carried out with MetaBAT2 ^110^, MaxBin 2.0 ^111^ and CONCOCT ^112^ within the wrapper metaWRAP v20200226^113^. Bins were evaluated using single copy marker genes as part of the classify workflow in checkM v1.1.3 ^114^, which provided completion and redundancy percentages for each bin. The best bins for each assembly were identified with metawrap’s refinement tool, including only bins that were estimated to be > 50% complete and < 10% redundant. As a measure of overall bin quality, quality scores (QS) were calculated using checkM statistics: bin QS = bin completeness – 5 x bin redundancy ^115^. Six bins with QS < 50% were targeted for further refinement via single assembly. One sample with high coverage for each bin and low coverage of all other bins was selected for single assembly with metaSPADES v3.14.0 ^116^. Reads from 92 randomly selected samples (and the six used for single assembly) were aligned back to each single assembly with bwa v0.7.17. Binning and refinement were carried out with metaWRAP as described above. We also inspected and manually refined bins using Anvi’o platform ^117^. The co-assembled and single assembled bins were then combined with bins generated in previous salmonid studies ^19, 31^. This list of 33 bins was then dereplicated at the approximate species-level (average nucleotide identity (ANI) > 95%) and a representative bin per ‘species’ identified with dRep v3.2.0 ^118^. These 16 representative bins form the final MAG catalogue used in all subsequent microbiota analyses. The MAG catalogue was taxonomically annotated with GTDB Tk v2.1.0 ^119^ using the GTDB r207 reference database ^120^. For each MAG, open reading frames were identified with Prodigal ^121^, single-copy core genes were identified with HMMER ^122^ and genes were annotated using the Pfam ^123^ and KEGG Orthologs databases ^124^, all within Anvi’o. The Illumina reads from each sample were then aligned back to the final MAG catalogue with bwa mem v0.7.17. Anvi’o v7.0 was then used to extract coverage and detection statistics for each sample’s alignment to each MAG.

For all downstream analyses (microbial community analyses, GWAS and MOFA), we defined a MAG as detected in a sample if ≥ 30% of the nucleotides in a MAG was covered by at least one read in that sample ^125^. MAGs with lower detection values were defined as not detected in that sample and their abundance was set to 0. One MAG (*Prevotella*) was assembled but was not detected in any sample after this filtering and is therefore not included in any analysis. We used Anvi’o ‘abundance’ values to represent normalised MAG abundance, defined as the mean coverage of a MAG in a sample divided by the overall mean coverage of all MAGs in a sample. For scaling in some downstream analyses, we then applied a log10 transformation to the abundance values (with a pseudocount of 0.001 for zero values).

We compared MAG abundances between samples and the negative controls. After filtering, no MAGs were detected in five of the seven negative controls. The remaining two negative controls had MAG01 present at detectable abundances after filtering (47–54 normalised mean coverage). No other MAGs were detected in these two controls. Since MAG01 was the most abundant MAG in our samples, this infrequent detection is likely due to cross-contamination during laboratory processing ^126^; Furthermore, MAG01 was not detected at all in the other five negative controls. Thus, we did not exclude MAG01 as a contaminant.

### 16S metabarcoding

Metabarcoding data of the 16S ribosomal RNA (rRNA gene) was generated for 140 gut epithelial scrapes. The sample collection was composed of gut content (60), gut tissue (60) and pellets of each feed type (10 extraction replicates for each feed type). First, DNA was extracted using the ZymoBIOMICS™ DNA/RNA MiniPrep kit (ZRC201962). For the gut content, 750 µL Shield™ (Zymo Research) was used as input for the extraction. For the gut tissue and feed types, approximately 150 mg were used as input for the extraction. Subsequently, the DNA sample extracts were screened for inhibition using qPCR ^127^ with bacterial 16S primers V3V4 ^128^. Moreover, the Ct values, as well as amplification plots, generated from the qPCR were used to estimate the number of tagged PCR cycles. Next, the samples were amplified using tagged PCR ^129^ with the bacterial 16S primers V3V4 designed with unique oligonucleotide tags. Thus, each amplified sample was tagged with a unique tag combination to identify potential tag-jumping errors ^130^. The gut content and feed types were given 25 cycles of tagged PCR, whereas the gut tissue were given 30 cycles of tagged PCR. The tagged PCR amplicons were then pooled according to gel band visibility and purified using SPRI beads ^131^. The purified amplicon pools were built into libraries using the PCR-free protocol TagSteady ^132^. Subsequently, the libraries were quantified using a 2100 BioAnalyzer Instrument™ (Agilent) and pooled equimolar. Ultimately, the library pool was sequenced on a Illumina MiSeq lane aiming for 107 Mb per sample of PE 350 bp data. Negative controls were included in all steps, from DNA extraction to library build, to detect potential microbial contaminants.

Sample demultiplexing and adapter removal was performed using AdapterRemoval v2.2.4 ^99^. Using the pipeline BAMSE v1.0 (https://github.com/anttonalberdi/bamse), primer clipping was performed by Cutadapt 2.10 ^133^, followed by read orientation, read quality trimming and read filtering. DADA2 v1.17.3 ^134^ was used for error learning and correction, dereplication, paired-end read merging and generation of amplicon sequence variants (ASVs). Chimeric ASVs were filtered by DADA2 and the remaining ASVs were assigned taxonomy at the genus level using the SILVA nonredundant SSU database v138. ASVs assigned to Eukaryotes (including mitochondria or chloroplast sequences) were removed, as were ASVs with a relative number of copies < 0.01% in each sample. Putative contaminant sequences from laboratory processing were identified and removed with decontam v1.8.0 ^135^.

To compare the 16S community composition between gut content, gut mucosa and feed pellet samples, we focused on ASVs present in at least 5 samples after quality control (Supplementary Figure 3). We generated a heatmap of ASV abundance using the R package pheatmap. ASV read counts per sample were normalised to abundance values using the centre-log-ratio transformation. We also generated a venn diagram using the R package ggVennDiagram, displaying the number of ASVs detected in and among each sample type.

### Gut metabolomics

Gut content samples were cryo-homogenised in 25% water, 25% methanol and 50% dichloromethane in a 1:15 sample:solvent ratio (w:v). Homogenisation was carried out using an OMNI Bead Ruptor 24, using liquid nitrogen to keep homogenised samples below 0°C to minimise degradation of metabolites during extraction. Homogenates were centrifuged at 20,000 g (0°C) and the polar phase from all samples was concentrated using SpeedVac (ThermoFisher Scientific) and resuspended in 200 µL 5% methanol. Four procedural blanks were included in homogenisation. A volume of 100 µl of all samples was collected into a Quality Control sample used for normalisation to enhance the detection of metabolites. Samples were measured on a nano-flow ultra-high pressure liquid chromatography–tandem high-resolution mass spectrometry analysis. Metabolites were detected using a Q Exactive™ HF Hybrid Quadrupole-Orbitrap™ Mass Spectrometer (ThermoFisher Scientific) operated in positive ion data-dependent acquisition mode ^136^.

ThermoFisher Scientific UHPLC-Orbitrap-MS/MS RAW files were converted into mzML files using Proteo Wizard ^137^. For an increased deciphering of molecular spectres, MZmine2 ^138, 139^ was applied for mass detection of MS1 and MS2 spectres, followed by chromatogram detection and deconvolution. Subsequently, detected isotopes and features were grouped according to a tolerance of mass-charges (5 ppm for m/z) and retention time (6 sec.) and the features were further aligned according to retention time and m/z. Lastly, only features with a MS2 spectrum were kept for further substructural analysis and *in silico* analysis. A molecular network was created using the feature-based molecular networking (FBMN) workflow on the Global Natural Product Social Molecular Networking (GNPS) platform ^140, 141^ combined with SIRIUS4 and CSI:FingerID ^142, 143^, as previously described for fish intestinal metabolomics ^144^. In brief, to enhance identification of unknown metabolites, unsupervised substructures were discovered using MS2LDA ^145, 146^ and MS2 spectra were annotated *in silico* using Network Annotation Propagation (NAP) ^147^. Furthermore, peptidic natural products (PNPs) were annotated *in silico*, using DEREPLICATOR ^148^. Chemical classes were retrieved for all GNPS library hits and *in silico* structures using ClassyFire ^149^. Finally, all structural annotations were combined within one network using MolNetEnhancer ^150^.

Metabolites detected in < 50% of all fish samples were filtered out to reduce zero inflation issues in the dataset, and samples with > 80% missing data were also excluded. Abundance data were then further processed in the R package MetaboDiff ^151^, where missing data was imputed for all metabolites detected in at least 60% of samples. Abundance data were normalised across samples to generate relative abundances. The PCA generated from these normalised abundances revealed strong batch effects, a common problem in metabolomic data ^152^. For the PCA and inclusion of metabolomics in the unsupervised MOFA analysis, we therefore used limma’s function removeBatchEffect ^153^ to regress out the metabolomic batch effect from the normalised data. PERMANOVA was performed on Euclidean distances of the normalised abundances, both before and after the removal of batch effects, using the adonis2 function and testing for associations with feed type, size class and batch (Supplementary Table 3).

### Fatty acid profiling

The fatty acid composition was analysed in both the experimental diets and fish muscle samples. Lipids from the samples were extracted by adding chloroform-methanol (2:1, v/v) and 19:0 methyl ester was added as an internal standard. After extraction of lipids, the samples were filtered, saponified and esterified using 12% BF_3_ in methanol. Fatty acid methyl esters were separated on a gas chromatograph and detected using a flame ionization detector, as described earlier ^154, 155^. The fatty acids were identified by retention time using standard mixtures of methyl esters (Nu-Chek, Elyian, USA), and quantified by the internal standard method.

For downstream analyses, the relative abundance (%) of each fatty acid was log10 transformed. MaAsLin2 ^156^ was used to identify differences in log-transformed fatty acid relative abundances between cestode detection, size class and feed types, while controlling for sampling date as a random effect. We classed associations as statistically significant with FDR (q value) < 0.05; however, we report all associations with FDR < 0.2 in Supplementary Table 2.

### Microbial community analysis

Alpha diversity from MAG abundance values was calculated using the Hill numbers framework with the R package hillR, using *q* = 1 to estimate the Shannon index ^157^. We used a linear model to test for statistically significant differences in the Shannon index by cestode detection, size class and feed type, while controlling for sequencing depth.

Nonmetric multidimensional scaling (NMDS) was performed using log10-transformed abundance values and Euclidean distance with the ordinate function in the R package phyloseq v1.40.0 ^158^. The NMDS stress value, which is a measure of the degree to which the distance between samples in the reduced dimensional space corresponds with the actual distance between samples (similar to a goodness-of-fit value), has been included in the figure legend of each NMDS plot. The k value (number of dimensions) is also given in each NMDS plot. PERMANOVAs were performed on the Euclidean distances using adonis2. NMDS using MAG presence/absence data and Jaccard distance was performed with the metaMDS function in vegan, and PERMANOVAs were performed on the Jaccard distances using adonis2. PERMANOVA models are provided in Supplementary Table 3.

We used MaAsLin2 ^156^ to identify differences in MAG abundance and MAG detection between cestode detection, size class and feed types, while controlling for sequencing depth as a fixed effect and sampling date as a random effect. For MAG abundance, MaAsLin2 was run with ‘normalisation = none’ and ‘transformation = log’ parameters, whereas for MAG detection, ‘transformation’ was also set to ‘none’. We classed associations as statistically significant with a false discovery rate (also called a q value) < 0.05; however, we report all associations with FDR < 0.2 in Supplementary Table 5.

For subsequent analyses, we used MAG abundance data (excluding MAGs detected in < 50 samples) and MAG detection data (i.e. presence/absence, excluding MAGs detected in < 10 samples). We also defined three additional ‘MAG detection’ variables: detection of any low-abundant mycoplasma MAG (i.e. MAG02–MAG05); detection of any *Photobacterium* MAG (MAG06–MAG07); and detection of any *Aliivibrio* MAG (MAG08–MAG10). Since MAG01 (*M. salmoninae*) was detected in all but 3 of the 392 metagenomes, we also defined a fourth ‘MAG detection’ binary variable, representing samples with ‘high’ vs ‘low’ abundance of MAG01 (> or ≤ 1.7 normalised abundance, respectively, based on the third quartile of the MAG01 abundance distribution).

### 16S profiles vs metagenomes

To confirm that our assembled MAGs captured most of the bacterial diversity in the gut content community, we compared our 16S ASV relative abundances to our MAG relative abundances. Since MAG and ASV taxonomy were assigned using different databases, we summed abundance across all gut content samples at the genus level (*Aliivibrio*, *Brevinema*, *Carnobacterium*, *Clostridium*, *Lactobacillus*, *Neilla*, *Photobacterium* and *Psychromonas*), family level (*Mycoplasmataceae*, *Vibrionaceae*) and order level (*Alteromonadales*). Total sum scaling was used to bring all abundance values within each dataset to a percentage (0-100%). Relative abundances were then compared visually against 1:1 hypothetical perfect concordance (Supplementary Figure 4). All taxa from the MAG catalogue are represented by at least one level in the plot, whereas all ASVs not in the above taxa lists were grouped as ‘Other’.

### Genome-wide association study

To identify genetic variation underlying phenotype differences, a Genome-Wide Association Study (GWAS) was performed with GEMMA ^159^, which uses linear mixed-models to test for genome-wide associations. A centred genotype relatedness matrix was included in the model to account for the high relatedness among individuals in the population. We used a ‘leave-one-chromosome-out’ strategy to prevent overcorrection of the model, where GEMMA was run on each chromosome separately and the current chromosome was not included in the relatedness matrix calculation. To account for wider population structure, principal components (PCs) were estimated with PCAngsd v20190125 ^160^ and the first five PCs were included as covariates in each GWAS run. Results were collated across chromosomes for the generation of qqplots, inflation factor (λ) and Manhattan plots in R. SNPs with p values < 5×10-8 were lassed as strongly associated with the phenotype of interest, while SNPs with p values < 1×10-5 were classified as moderately associated.

For size investigations, gutted weight was transformed with the inverse normal transformation to approximate a normal distribution for linear regression in the model. Feed type (converted to a binary variable) was included as a covariate. GEMMA was also run with feed type as a phenotype, with gutted weight (untransformed) as a covariate. Detection of cestode (binary presence/absence) was also tested, including gutted weight (untransformed) and feed type as covariates. For microbiome (m)GWAS, microbiome phenotypes were included as binary variables (MAG detection) and INT-transformed (MAG abundance, alpha diversity) or untransformed (ordination dimensions) continuous variables. Feed type, gutted weight (untransformed) and cestode presence/absence were included as covariates along with PCs1-5 for each mGWAS to account for wider population structure.

Only one microbiome phenotype was associated with host genetic variation in the mGWAS, specifically MAG05 presence or absence and 14 SNPs in a 1.8-Mbp region of chr5. The locations and annotations of candidate associated SNPs in the genome annotation file were identified with BEDtools v2.30.0 ^161^, extracting gene and mRNA annotations overlapping (‘bedtools intersect’) with the coordinates of each SNP. For SNPs located within a gene or mRNA transcript, further annotation information, e.g. Gene Ontology (GO) terms, were accessed using the R package AnnotationDBI v1.58 to query the Atlantic salmon annotation stored in the R package AnnotationHub v3.4 (organism AH100820, last updated 21-04-2022). GO term descriptions were accessed using the R package GO.db v3.15. The coordinate of each candidate SNP was also entered into the Ensembl ‘Variant Effect Predictor’ tool (https://www.ensembl.org/Tools/VEP), to determine whether the SNP (relative to the reference genome) resulted in a synonymous or nonsynonymous mutation in a coding region, or if it fell in an intron or noncoding region.

To investigate the effect of these SNPs on gene expression, we used the genotype probabilities to define genotypes for each individual at each of the 14 significant positions, using a genotype probability > 0.8 for categorise an individual as major/major (MM), major/minor (Mm) or minor/minor (mm). We excluded individuals with genotype probabilities ≤ 0.8 for all three genotypes. For SNPs located in genes that were sequenced in our transcriptomic dataset, we compared expression of those genes (post VST normalisation) between individuals with the major/major, major/minor and minor/minor genotypes, and between fish with MAG05 present or absent. Statistically significant differences in gene expression (p < 0.05) were identified using Wilcoxon rank sum tests implemented in the R package ggsignif ^162^. To identify more widespread effects of these SNPs on gene expression, we performed differential expression analysis using DESeq, as described in ‘Host gut transcriptomics’, using the subset of 36 genes that were located within 1 Mbp in either direction of the chr 5 SNP peak. Two of the 14 SNPs had the same genotype in all individuals included in the transcriptomic analysis and were therefore excluded from this analysis. We ran three different models in DESeq1: 1) the 12 SNP genotypes, 2) MAG05 presence/absence and 3) the 12 SNP genotypes and MAG05 presence/absence. In all three models, we also included feed type, size class and cestode presence/absence as covariates.

### Multi-omics factor analysis

Multi-omics factor analysis (MOFA) was used to integrate the metagenome, metabolome and transcriptome datasets, using the R package MOFA2 ^38^. For the metagenome dataset, we included presence/absence data for all MAGs detected in > 10 and < 388 samples (11 MAGs) as well as the four additional MAG detection variables (any low-abundant *Mycoplasma* MAG, any *Photobacterium* MAG, any *Aliivibrio* MAG and ‘high’ vs ‘low’ detection of MAG01). For the metabolome dataset, we included the 500 most variable metabolite abundances (normalised as described in ‘Gut metabolomics’). For the host transcriptome dataset, we included the 500 most variable gene expression abundances (following DegNorm, DESeq2 and vst processing steps, as described in ‘Host gut transcriptomics’). We trained the MOFA model with 15 factors, using Gaussian (metabolome and transcriptome) and Bernoulli (MAG detection) distributions, scaling each dataset to have similar variances and otherwise using default values. To identify associations that were independent of feed type, we separated samples into two groups by feed type and trained a MOFA model on each, using the same parameters as described for the full model.

After each model was trained, we used the function correlate_factors_with_covariates in the MOFA2 package to identify factors that were significantly correlated (alpha = 0.05) with cestode detection, size class and feed type, using Pearson correlation coefficients and associated p-values (adjusted for multiple hypothesis testing). We focused on factors that significantly (adjusted p < 0.05) correlated with cestode detection and/or size class, and those that explained >1% of the variation in at least two -omic datasets. However, the full results of the MOFA models can be found on the study’s GitHub (https://github.com/jcbrealey/HoloFish_multiomics). We report all features (MAG variables, metabolites or genes) with absolute weights > 0.2 as significantly contributing to these factors. We compared the results of the full model to each model based on feed type, and labelled a feature association as ‘robust’ if it significantly contributed to a factor in both feed type models that was associated with the same fish variable (cestode detection or size class) in the full model. For visualisation, MAG features were annotated with their assigned genus, metabolite features were annotated by metabolite class and gene features were annotated by key words in their associated GO terms or gene names (script available at https://github.com/jcbrealey/HoloFish_multiomics).

### MAG phylogenetics and comparative genomics

Phylogenetics and comparasite genomics for the *Mycoplasma* and *Vibrionaceae* MAGs were performed with the Anvi’o platform ^117^. Related *Mycoplasmataceae* and *Vibrionaceae* reference species genomes (Supplementary Table 13) were annotated with Anvi’o using the Pfam ^123^ and KEGG Orthologs databases ^124^, as described for each MAG above. Pangenome profiles were generated in Anvi’o for the *Mycoplasmataceae* and *Vibrionaceae* genomes separately, including the identification of gene clusters by alignment with DIAMOND ^163^ in ‘fast’ mode, elimination of weak alignments with the minbit heuristic ^164^ and clustering with MCL ^165^. Presence and completeness of KEGG metabolic pathways were estimated with the anvi-estimate-metabolism function. Amino acid sequences of single copy bacterial ribosomal gene clusters (the Bacteria_71 collection in Anvi’o) were extracted, concatenated and constructed into a phylogenomic tree using the Anvi’o implementation of FastTree ^166^. FigTree v1.4.4 (http://tree.bio.ed.ac.uk/software/figtree/) was used to visualise the resulting tree and reroot it using the outgroup, *Ureaplasma* for *Mycoplasmataceae* and *Vibrio* for *Vibrionaceae*. Pairwise ANI was estimated using the Anvi’o implementation of PyANI ^167^.

For functional analysis, a literature search was performed to compile a list of genes known to be important for colonisation, survival and virulence in terrestrial and marine hosts of *Mycoplasmataceae* and *Vibrionaceae* ^39, 43, 47, 50, 168–170^. Where possible, KEGG pathways and KEGG orthologue and Pfam annotation of gene clusters were searched for these gene names or gene symbols. For genes that were less well-characterised (e.g. adhesins lacking formal names), we downloaded reference sequences of these genes and compared them to the sequences of our pangenome gene clusters, using blastp v2.13 ^171^. A gene was defined as present in our pangenome if it had a corresponding KEGG or Pfam annotation, or if ≥ 50% of the reference sequence was aligned to a gene cluster with an E value < 0.01. A complete list of genes and accessions are reported in Supplementary Table 14. Only those present in at least one MAG are shown in Figure 5.

## Data availability

Raw metagenomic, metabarcoding, genomic and transcriptomic sequencing data are available on the European Nucleotide archive (ENA) under project PRJEB64334. Individual metadata are available as BioSamples under the same project. All sample and run ENA accessions are provided in Supplementary Table 16. Assembled MAG sequences are available as analysis files under the same ENA project, with accessions provided in Supplementary Table 1. Metabolomic data are available on MassIVE under the dataset identifier XXXX. Fatty acid composition data and all analysis files and scripts are available at https://github.com/jcbrealey/HoloFish_multiomics.

## Author contributions

JCB, LM, HS, KK, MTPG, MDM and MTL designed the study. Project management was undertaken by MTPG, MDM and MTL. Sampling was performed by MK, JAR, EF, AB, MTPG, MDM and MTL. Data was generated by MK, JAR, LSB, LAL, MH, EF, LSM and AB while data analysis was performed by JCB, MK, JAR, SBH, LAL, EF, LSM and AB. JCB predominately wrote the manuscript, with contributions on methods text by MK, JAR, LSB, MH, EF, LSM and AB. All authors edited and approved the final manuscript.

## Acknowledgements

We are thankful to Maria Limborg Karm, Filipe Garrett Vieira, Mikkel Holger Strander Sinding, Martin Nielsen and Cathrine Kalgraff for help with sampling salmon tissues in the field. We also thank Sen Li for assistance with the -omic data processing pipelines, Simen Rød Sandve and his team at Norwegian University of Life Sciences for early access to the salmon reference genome and Shyam Gopalakrishnan for valuable advice on performing the GWAS analysis.

## Funding

The work was funded by the FHF-Norwegian Seafood Research Fund (“HoloFish,” grant no. 901436), the Danish National Research Foundation (CEH-DNRF143), the European Union’s Horizon 2020 Action (“HoloFood”, grant no. 817729), the European Union’s Horizon 2020 ERA-Net Cofund project BlueBio (“ImprovAFish”, grant no. 311913) and the Carlsberg foundation’s Semper Ardens accelerate grant (“EpoEvo”, grant no. CF21-0356).

## Conflicts of interest

The authors declare a conflict of interest. Harald Sveier is employed at Lerøy Seafood Group, who produce, market and sell farmed Atlantic salmon including the fish used in the current investigation. Furthermore, Lerøy Seafood Group provided parts of the funding for this study. All other authors declare no competing interests.

## Supplementary Material

### Supplementary Figures

- Supplementary Figure 1. Ordination (NMDS) of metagenomes, based on MAG abundance and MAG presence/absence.
- Supplementary Figure 2. Alpha diversity (Hill’s shannon index) of metagenomes, based on MAG abundance, stratified by cestode detection and size class.
- Supplementary Figure 3. Microbiota community composition of salmon gut content compared to salmon gut mucosa scrapes and feed pellet, determined by 16S metabarcoding.
- Supplementary Figure 4. Comparison of microbial relative abundances in salmon gut content samples captured by MAG assembly versus 16S profiling.
- Supplementary Figure 5. Manhattan plots of all GWAS runs.
- Supplementary Figure 6. Ordination (PCA) of salmon transcriptomes.
- Supplementary Figure 7. Ordination (PCA) of gut metabolomes.
- Supplementary Figure 8. MOFA results for factors of interest in salmon fed Feed1.
- Supplementary Figure 9. MOFA results for factors of interest in salmon fed Feed2.

### Supplementary Tables

- Supplementary Table 1. MAG quality statistics and other information.
- Supplementary Table 2. Associations between fish phenotypes and fatty acid relative abundances in muscle.
- Supplementary Table 3. PERMANOVAs of variation in salmon gut metagenomes, transcriptomes and metabolomes.
- Supplementary Table 4. Linear models of metagenome alpha diversity differences among fish phenotypes.
- Supplementary Table 5. Associations between fish phenotypes and metagenomes, based on MAG abundance and MAG presence/absence.
- Supplementary Table 6. Summary of GWAS results for metagenome phenotypes and fish phenotypes.
- Supplementary Table 7. Information and annotation of SNPs associated with MAG05 on chromosome 5 of the salmon genome.
- Supplementary Table 8. MOFA summaries for each model.
- Supplementary Table 9. MOFA metagenome feature weights and annotations for factors of interest and significant features in the combined model.
- Supplementary Table 10. MOFA metabolome feature weights and annotations for factors of interest and significant features in the combined model.
- Supplementary Table 11. MOFA transcriptome feature weights and annotations for factors of interest and significant features in the combined model.
- Supplementary Table 12. ANI percentages among (A) *Mycoplasma* and (B) *Aliivibrio* and *Photobacterium* MAGs.
- Supplementary Table 13. Bacterial reference genomes used for MAG comparative genomics.
- Supplementary Table 14. Bacterial genes targeted in MAG comparative genomics and Figure 5.
- Supplementary Table 15. Differentially expressed host genes on chromosome 5 among genotypes and MAG detection, as identified in the GWAS analysis.
- Supplementary Table 16. Individual salmon metadata, samples and ENA accessions.

## Notes

### Competing Interest Statement

Conflicts of interest
The authors declare a conflict of interest. Harald Sveier is employed at Leroy Seafood Group, who produce, market and sell farmed Atlantic salmon including the fish used in the current investigation. Furthermore, Leroy Seafood Group provided parts of the funding for this study. All other authors declare no competing interests.

## References

1. Zheng, D., Liwinski, T. & Elinav, E. Interaction between microbiota and immunity in health and disease. Cell Res. 30, 492–506 (2020).

2. Kelly, C. & Salinas, I. Under pressure: Interactions between commensal microbiota and the teleost immune system. Front. Immunol. 8, 559 (2017).

3. Gensollen, T., Iyer, S. S., Kasper, D. L. & Blumberg, R. S. How colonization by microbiota in early life shapes the immune system. Science 352, 539–544 (2016).

4. Hacquard, S. et al. Microbiota and Host Nutrition across Plant and Animal Kingdoms. Cell Host Microbe 17, 603–616 (2015).

5. Qin, Y. et al. Combined effects of host genetics and diet on human gut microbiota and incident disease in a single population cohort. Nat. Genet. 54, 134–142 (2022).

6. Lopera-Maya, E. A. et al. Effect of host genetics on the gut microbiome in 7,738 participants of the Dutch Microbiome Project. Nat. Genet. 54, 143–151 (2022).

7. Sevellec, M., Laporte, M., Bernatchez, A., Derome, N. & Bernatchez, L. Evidence for host effect on the intestinal microbiota of whitefish (*Coregonus* sp.) species pairs and their hybrids. Ecol. Evol. 9, 11762–11774 (2019).

8. Fietz, K. et al. Mind the gut: Genomic insights to population divergence and gut microbial composition of two marine keystone species. Microbiome 6, 82 (2018).

9. Richards, A. L. et al. Gut microbiota has a widespread and modifiable effect on host gene regulation. mSystems 4, e00323–18 (2017).

10. Nyholm, L. et al. Holo-Omics: Integrated Host-Microbiota Multi-omics for Basic and Applied Biological Research. iScience 23, 101414 (2020).

11. Reynolds, L. A., Finlay, B. B. & Maizels, R. M. Cohabitation in the Intestine: Interactions among Helminth Parasites, Bacterial Microbiota, and Host Immunity. J Immunol 195, 4059–4066 (2015).

12. Williams, A. R. et al. Emerging interactions between diet, gastrointestinal helminth infection, and the gut microbiota in livestock. BMC Vet. Res. 17, 62 (2021).

13. Hahn, M. A. et al. Host phenotype and microbiome vary with infection status, parasite genotype, and parasite microbiome composition. Mol. Ecol. (2022) doi:10.1111/mec.16344.

14. Gaulke, C. A. et al. A longitudinal assessment of host-microbe-parasite interactions resolves the zebrafish gut microbiome’s link to *Pseudocapillaria tomentosa* infection and pathology. Microbiome 7, 10 (2019).

15. White, E. C. et al. Manipulation of host and parasite microbiotas: Survival strategies during chronic nematode infection. Science Advances 4, eaap7399 (2018).

16. Plieskatt, J. L. et al. Infection with the carcinogenic liver fluke *Opisthorchis viverrini* modifies intestinal and biliary microbiome. FASEB J. 27, 4572–4584 (2013).

17. Jaenike, J., Unckless, R., Cockburn, S. N., Boelio, L. M. & Perlman, S. J. Adaptation via symbiosis: Recent spread of a drosophila defensive symbiont. Science 329, 212–215 (2010).

18. Jorge, F., Dheilly, N. M. & Poulin, R. Persistence of a Core Microbiome Through the Ontogeny of a Multi-Host Parasite. Front. Microbiol. 11, 954 (2020).

19. Brealey Jaelle C., et al. Microbiome ‘Inception’: an Intestinal Cestode Shapes a Hierarchy of Microbial Communities Nested within the Host. mBio 13, e00679–22 (2022).

20. Kashinskaya, E. N. et al. Trophic diversification and parasitic invasion as ecological niche modulators for gut microbiota of whitefish. Front. Microbiol. 14, 1090899 (2023).

21. Landmann, F., Voronin, D., Sullivan, W. & Taylor, M. J. Anti-filarial activity of antibiotic therapy is due to extensive apoptosis after *Wolbachia* depletion from filarial nematodes. PLoS Pathog. 7, e1002351 (2011).

22. Vaughan, J. A., Tkach, V. V. & Greiman, S. E. Neorickettsial Endosymbionts of the Digenea: Diversity, Transmission and Distribution. vol. 79 253–297 (Elsevier, 2012).

23. Dessi, D., Rappelli, P., Diaz, N., Cappuccinelli, P. & Fiori, P. L. *Mycoplasma hominis* and *Trichomonas vaginalis*: A unique case of symbiotic relationship between two obligate human parasites. Front. Biosci. 11, 2028–2034 (2006).

24. Deenonpoe, R. et al. The carcinogenic liver fluke *Opisthorchis viverrini* is a reservoir for species of *Helicobacter*. Asian Pac. J. Cancer Prev. 16, 1751–1758 (2015).

25. Zilber-Rosenberg, I. & Rosenberg, E. Role of microorganisms in the evolution of animals and plants: the hologenome theory of evolution. FEMS Microbiol. Rev. 32, 723– 735 (2008).

26. Theis, K. R. et al. Getting the Hologenome Concept Right: an Eco-Evolutionary Framework for Hosts and Their Microbiomes. mSystems 1, e00028–16 (2016).

27. Alberdi, A., Andersen, S. B., Limborg, M. T., Dunn, R. R. & Gilbert, M. T. P. Disentangling host-microbiota complexity through hologenomics. Nat. Rev. Genet. (2021) doi:10.1038/s41576-021-00421-0.

28. Limborg, M. T. et al. Applied Hologenomics: Feasibility and Potential in Aquaculture. Trends Biotechnol. 36, 252–264 (2018).

29. Bristow, G. A. & Berland, B. The effect of long term, low level *Eubothrium* sp. (Cestoda: Pseudophyllidea) infection on growth in farmed salmon (Salmo salar L.). Aquaculture 98, 325–330 (1991).

30. Saksvik, M., Nilsen, F., Nylund, A. & Berland, B. Effect of marine *Eubothrium* sp. (Cestoda: Pseudophyllidea) on the growth of Atlantic salmon, *Salmo salar* L. J. Fish Dis. 24, 111–119 (2001).

31. Rasmussen, J. A. et al. Genome-resolved metagenomics suggests a mutualistic relationship between *Mycoplasma* and salmonid hosts. Communications Biology 4, 579 (2021).

32. Rasmussen, J. A. et al. Co-diversification of an intestinal *Mycoplasma* and its salmonid host. ISME J. (2023) doi:10.1038/s41396-023-01379-z.

33. Bozzi, D. et al. Salmon gut microbiota correlates with disease infection status: potential for monitoring health in farmed animals. Animal Microbiome 3, 30 (2021).

34. Holben, W. E., Williams, P., Saarinen, M., Särkilahti, L. K. & Apajalahti, J. H. A. Phylogenetic Analysis of Intestinal Microflora Indicates a Novel Mycoplasma Phylotype in Farmed and Wild Salmon. Microb. Ecol. 44, 175–185 (2002).

35. Dehler, C. E., Secombes, C. J. & Martin, S. A. M. Seawater transfer alters the intestinal microbiota profiles of Atlantic salmon (*Salmo salar* L.). Sci. Rep. 7, 13877 (2017).

36. Llewellyn, M. S. et al. The biogeography of the atlantic salmon (*Salmo salar*) gut microbiome. ISME J. 10, 1280–1284 (2016).

37. Li, Y. et al. Differential response of digesta- and mucosa-associated intestinal microbiota to dietary insect meal during the seawater phase of Atlantic salmon. Anim Microbiome 3, 8 (2021).

38. Argelaguet, R. et al. Multi-Omics Factor Analysis-a framework for unsupervised integration of multi-omics data sets. Mol. Syst. Biol. 14, e8124 (2018).

39. Yiwen, C., Yueyue, W., Lianmei, Q., Cuiming, Z. & Xiaoxing, Y. Infection strategies of mycoplasmas: Unraveling the panoply of virulence factors. Virulence 12, 788–817 (2021).

40. Blötz, C. & Stülke, J. Glycerol metabolism and its implication in virulence in *Mycoplasma*. FEMS Microbiol. Rev. 41, 640–652 (2017).

41. Imanishi, I. et al. Exfoliative toxin E, a new *Staphylococcus aureus* virulence factor with host-specific activity. Sci. Rep. 9, 16336 (2019).

42. Fernandez, R. C. & Weiss, A. A. Cloning and sequencing of a *Bordetella pertussis* serum resistance locus. Infect. Immun. 62, 4727–4738 (1994).

43. Metsugi, S. et al. Sequence analysis of the gliding protein Gli349 in *Mycoplasma mobile*. Biophysics (Nagoya-shi) 1, 33–43 (2005).

44. Onarheim, A. M., Wiik, R., Burghardt, J. & Stackebrandt, E. Characterization and Identification of Two Vibrio Species Indigenous to the Intestine of Fish in Cold Sea Water; Description of *Vibrio iliopiscarius* sp. nov. Syst. Appl. Microbiol. 17, 370–379 (1994).

45. Budsberg, K. J., Wimpee, C. F. & Braddock, J. F. Isolation and identification of *Photobacterium phosphoreum* from an unexpected niche: migrating salmon. Appl. Environ. Microbiol. 69, 6938–6942 (2003).

46. Ast, J. C. & Dunlap, P. V. Phylogenetic resolution and habitat specificity of members of the *Photobacterium phosphoreum* species group. Environ. Microbiol. 7, 1641–1654 (2005).

47. Urbanczyk, H., Ast, J. C. & Dunlap, P. V. Phylogeny, genomics, and symbiosis of *Photobacterium*. FEMS Microbiol. Rev. 35, 324–342 (2011).

48. Visick, K. L., Stabb, E. V. & Ruby, E. G. A lasting symbiosis: how *Vibrio fischeri* finds a squid partner and persists within its natural host. Nat. Rev. Microbiol. 19, 654–665 (2021).

49. Riemann, L. & Azam, F. Widespread N-acetyl-D-glucosamine uptake among pelagic marine bacteria and its ecological implications. Appl. Environ. Microbiol. 68, 5554–5562 (2002).

50. Hjerde, E. et al. The genome sequence of the fish pathogen *Aliivibrio salmonicida* strain LFI1238 shows extensive evidence of gene decay. BMC Genomics 9, 616 (2008).

51. Pellock, S. J. & Redinbo, M. R. Glucuronides in the gut: Sugar-driven symbioses between microbe and host. J. Biol. Chem. 292, 8569–8576 (2017).

52. Jin, Y. et al. Atlantic salmon raised with diets low in long-chain polyunsaturated n-3 fatty acids in freshwater have a *Mycoplasma*-dominated gut microbiota at sea. Aquac. Environ. Interact. 11, 31–39 (2019).

53. McKenney, E. A. et al. Alteration of the rat cecal microbiome during colonization with the helminth *Hymenolepis diminuta*. Gut Microbes 6, 182–193 (2015).

54. Aivelo, T. & Norberg, A. Parasite–microbiota interactions potentially affect intestinal communities in wild mammals. J. Anim. Ecol. 87, 438–447 (2018).

55. Sheehy, L. et al. A parasitic nematode induces dysbiosis in susceptible but not resistant gastropod hosts. Microbiologyopen 12, (2023).

56. Liang, L., Ai, L., Qian, J., Fang, J.-Y. & Xu, J. Long noncoding RNA expression profiles in gut tissues constitute molecular signatures that reflect the types of microbes. Sci. Rep. 5, 11763 (2015).

57. Li, Z. et al. The long noncoding RNA *THRIL* regulates TNFα expression through its interaction with hnRNPL. Proc. Natl. Acad. Sci. U. S. A. 111, 1002–1007 (2014).

58. Carpenter, S. et al. A long noncoding RNA mediates both activation and repression of immune response genes. Science 341, 789–792 (2013).

59. Boltaña, S., Valenzuela-Miranda, D., Aguilar, A., Mackenzie, S. & Gallardo-Escárate, C. Long noncoding RNAs (lncRNAs) dynamics evidence immunomodulation during ISAV-Infected Atlantic salmon (*Salmo salar*). Sci. Rep. 6, 22698 (2016).

60. Ling, F. et al. The gut microbiota response to helminth infection depends on host sex and genotype. ISME J. 14, 1141–1153 (2020).

61. Romano, N. et al. Bile acid metabolism in fish: disturbances caused by fishmeal alternatives and some mitigating effects from dietary bile inclusions. Rev. Aquac. (2020) doi:10.1111/raq.12410.

62. Guzior, D. V. & Quinn, R. A. Review: microbial transformations of human bile acids. Microbiome 9, 140 (2021).

63. Wen, J. et al. Fxr signaling and microbial metabolism of bile salts in the zebrafish intestine. Sci Adv 7, (2021).

64. Gilbert, B. M. et al. You are how you eat: differences in trophic position of two parasite species infecting a single host according to stable isotopes. Parasitol. Res. 119, 1393– 1400 (2020).

65. Tkachuck, R. D. & MacInnis, A. J. The effect of bile salts on the carbohydrate metabolism of two species of hymenolepidid cestodes. Comparative Biochemistry and Physiology Part B: Comparative Biochemistry 40, 993–1003 (1971).

66. Kortner, T. M., Björkhem, I., Krasnov, A., Timmerhaus, G. & Krogdahl, Å. Dietary cholesterol supplementation to a plant-based diet suppresses the complete pathway of cholesterol synthesis and induces bile acid production in Atlantic salmon (*Salmo salar* L.). Br. J. Nutr. 111, 2089–2103 (2014).

67. Peñaloza, C., Hamilton, A., Guy, D. R., Bishop, S. C. & Houston, R. D. A SNP in the 5’ flanking region of the myostatin-1b gene is associated with harvest traits in Atlantic salmon (*Salmo salar*). BMC Genet. 14, 112 (2013).

68. Tsai, H. Y., Hamilton, A., Guy, D. R. & Houston, R. D. Single nucleotide polymorphisms in the insulin-like growth factor 1 (IGF1) gene are associated with growth-related traits in farmed Atlantic salmon. Anim. Genet. 45, 709–715 (2014).

69. Thodesen, J., Grisdale-Helland, B., Helland, S. J. & Gjerde, B. Feed intake, growth and feed utilization of offspring from wild and selected Atlantic salmon (*Salmo salar*). Aquaculture 180, 237–246 (1999).

70. Harvey, A. C. et al. Plasticity in growth of farmed and wild Atlantic salmon: is the increased growth rate of farmed salmon caused by evolutionary adaptations to the commercial diet? BMC Evol. Biol. 16, 264 (2016).

71. Solberg, M. F., Skaala, Ø., Nilsen, F. & Glover, K. A. Does domestication cause changes in growth reaction norms? A study of farmed, wild and hybrid Atlantic salmon families exposed to environmental stress. PLoS One 8, e54469 (2013).

72. Solberg, M. F., Kvamme, B. O., Nilsen, F. & Glover, K. A. Effects of environmental stress on mRNA expression levels of seven genes related to oxidative stress and growth in Atlantic salmon *Salmo salar* L. of farmed, hybrid and wild origin. BMC Res. Notes 5, 672 (2012).

73. Madaro, A. et al. Stress in Atlantic salmon: response to unpredictable chronic stress. J. Exp. Biol. 218, 2538–2550 (2015).

74. Fjelldal, P. G., Hansen, T. J. & Karlsen, Ø. Effects of laboratory salmon louse infection on osmoregulation, growth and survival in Atlantic salmon. Conserv Physiol 8, coaa023 (2020).

75. Gjøen, H. M. & Bentsen, H. B. Past, present, and future of genetic improvement in salmon aquaculture. ICES J. Mar. Sci. 54, 1009–1014 (1997).

76. Baranski, M., Moen, T. & Våge, D. I. Mapping of quantitative trait loci for flesh colour and growth traits in Atlantic salmon (*Salmo salar*). Genet. Sel. Evol. 42, 17 (2010).

77. Tsai, H. Y. et al. The genetic architecture of growth and fillet traits in farmed Atlantic salmon (*Salmo salar*). BMC Genet. 16, 51 (2015).

78. Thépot, V. et al. Dietary inclusion of the red seaweed *Asparagopsis taxiformis* boosts production, stimulates immune response and modulates gut microbiota in Atlantic salmon, Salmo salar. Aquaculture 546, 737286 (2022).

79. Hartviksen, M. et al. Alternative dietary protein sources for Atlantic salmon (*Salmo salar* L.) effect on intestinal microbiota, intestinal and liver histology and growth. Aquacult. Nutr. 20, 381–398 (2014).

80. Leeper, A. et al. Torula yeast in the diet of Atlantic salmon *Salmo salar* and the impact on growth performance and gut microbiome. Sci. Rep. 12, 567 (2022).

81. Bledsoe, J. W., Pietrak, M. R., Burr, G. S., Peterson, B. C. & Small, B. C. Functional feeds marginally alter immune expression and microbiota of Atlantic salmon (*Salmo salar*) gut, gill, and skin mucosa though evidence of tissue-specific signatures and host-microbe coadaptation remain. Anim Microbiome 4, 20 (2022).

82. Wang, C., Sun, G., Li, S., Li, X. & Liu, Y. Intestinal microbiota of healthy and unhealthy Atlantic salmon *Salmo salar* L. in a recirculating aquaculture system. Journal of Oceanology and Limnology 36, 414–426 (2018).

83. Egidius, E., Wiik, R. & Andersen, K. *Vibrio salmonicida* sp. nov., a new fish pathogen. Int. J. Syst. Bacteriol. 36, 518–520 (1986).

84. Young, K. M. et al. Bacterial-binding activity and plasma concentration of ladderlectin in rainbow trout (*Oncorhynchus mykiss*). Fish Shellfish Immunol. 23, 305–315 (2007).

85. Russell, S., Young, K. M., Smith, S., Hayes, M. A. & Lumsden, J. S. Cloning, binding properties, and tissue localization of rainbow trout (*Oncorhynchus mykiss*) ladderlectin. Fish & Shellfish Immunology 24, 669–683 (2008).

86. Bridle, A., Nosworthy, E., Polinski, M. & Nowak, B. Evidence of an antimicrobial-immunomodulatory role of Atlantic salmon cathelicidins during infection with *Yersinia ruckeri*. PLoS One 6, e23417 (2011).

87. Cham, B. E. Importance of apolipoproteins in lipid metabolism. Chem. Biol. Interact. 20, 263–277 (1978).

88. Concha, M. I., Molina, S., Oyarzún, C., Villanueva, J. & Amthauer, R. Local expression of apolipoprotein A-I gene and a possible role for HDL in primary defence in the carp skin. Fish Shellfish Immunol. 14, 259–273 (2003).

89. Villarroel, F., Bastías, A., Casado, A., Amthauer, R. & Concha, M. I. Apolipoprotein A-I, an antimicrobial protein in *Oncorhynchus mykiss*: evaluation of its expression in primary defence barriers and plasma levels in sick and healthy fish. Fish Shellfish Immunol. 23, 197–209 (2007).

90. Park, K. C., Osborne, J. A., Tsoi, S. C. M., Brown, L. L. & Johnson, S. C. Expressed sequence tags analysis of Atlantic halibut (*Hippoglossus hippoglossus*) liver, kidney and spleen tissues following vaccination against *Vibrio anguillarum* and *Aeromonas salmonicida*. Fish Shellfish Immunol. 18, 393–415 (2005).

91. Easy, R. H. & Ross, N. W. Changes in Atlantic salmon (*Salmo salar*) epidermal mucus protein composition profiles following infection with sea lice (*Lepeophtheirus salmonis*). Comp. Biochem. Physiol. Part D Genomics Proteomics 4, 159–167 (2009).

92. Wang, J. et al. Microbiota in intestinal digesta of Atlantic salmon (*Salmo salar*), observed from late freshwater stage until one year in seawater, and effects of functional ingredients: a case study from a commercial sized research site in the Arctic region. Anim Microbiome 3, 14 (2021).

93. Harpaz, S. l-Carnitine and its attributed functions in fish culture and nutrition—a review. Aquaculture 249, 3–21 (2005).

94. Tocher, D. R. Metabolism and Functions of Lipids and Fatty Acids in Teleost Fish. Rev. Fish. Sci. 11, 107–184 (2003).

95. Shulgina, N. S., Churova, M. V., Murzina, S. A., Krupnova, M. Y. & Nemova, N. N. The Effect of Continuous Light on Growth and Muscle-Specific Gene Expression in Atlantic Salmon (*Salmo salar* L.) Yearlings. Life 11, (2021).

96. Bentley, A. A. & Adams, J. C. The evolution of thrombospondins and their ligand-binding activities. Mol. Biol. Evol. 27, 2187–2197 (2010).

97. Carøe, C. et al. Single-tube library preparation for degraded DNA. Methods Ecol. Evol. 9, 410–419 (2018).

98. Mak, S. S. T. et al. Comparative performance of the BGISEQ-500 vs Illumina HiSeq2500 sequencing platforms for palaeogenomic sequencing. Gigascience 6, gix049 (2017).

99. Schubert, M., Lindgreen, S. & Orlando, L. AdapterRemoval v2: Rapid adapter trimming, identification, and read merging. BMC Res. Notes 9, 88 (2016).

100. Shen, W., Le, S., Li, Y. & Hu, F. SeqKit: A Cross-Platform and Ultrafast Toolkit for FASTA/Q File Manipulation. PLoS One 11, e0163962 (2016).

101. Li, H. Aligning sequence reads, clone sequences and assembly contigs with BWA-MEM. arXiv preprint arXiv:1303.3997 (2013).

102. Li, H. et al. The Sequence Alignment/Map format and SAMtools. Bioinformatics 25, 2078–2079 (2009).

103. McKenna, A. et al. The Genome Analysis Toolkit: a MapReduce framework for analyzing next-generation DNA sequencing data. Genome Res. 20, 1297–1303 (2010).

104. Korneliussen, T. S., Albrechtsen, A. & Nielsen, R. ANGSD: Analysis of Next Generation Sequencing Data. BMC Bioinformatics 15, 356 (2014).

105. Browning, B. L., Zhou, Y. & Browning, S. R. A One-Penny Imputed Genome from Next-Generation Reference Panels. Am. J. Hum. Genet. 103, 338–348 (2018).

106. Dobin, A. et al. STAR: ultrafast universal RNA-seq aligner. Bioinformatics 29, 15–21 (2013).

107. Xiong, B., Yang, Y., Fineis, F. R. & Wang, J.-P. DegNorm: normalization of generalized transcript degradation improves accuracy in RNA-seq analysis. Genome Biol. 20, 75 (2019).

108. Love, M. I., Huber, W. & Anders, S. Moderated estimation of fold change and dispersion for RNA-seq data with DESeq2. Genome Biol. 15, 550 (2014).

109. Li, D., Liu, C.-M., Luo, R., Sadakane, K. & Lam, T.-W. MEGAHIT: an ultra-fast single-node solution for large and complex metagenomics assembly via succinct de Bruijn graph. Bioinformatics 31, 1674–1676 (2015).

110. Kang, D. D., Froula, J., Egan, R. & Wang, Z. MetaBAT, an efficient tool for accurately reconstructing single genomes from complex microbial communities. PeerJ 3, e1165 (2015).

111. Wu, Y.-W., Simmons, B. A. & Singer, S. W. MaxBin 2.0: an string-nameomated binning algorithm to recover genomes from multiple metagenomic datasets. Bioinformatics 32, 605–607 (2016).

112. Alneberg, J. et al. Binning metagenomic contigs by coverage and composition. Nat. Methods 11, 1144–1146 (2014).

113. Uritskiy, G. V., DiRuggiero, J. & Taylor, J. MetaWRAP-a flexible pipeline for genome-resolved metagenomic data analysis. Microbiome 6, 158 (2018).

114. Parks, D. H., Imelfort, M., Skennerton, C. T., Hugenholtz, P. & Tyson, G. W. CheckM: assessing the quality of microbial genomes recovered from isolates, single cells, and metagenomes. Genome Res. 25, 1043–1055 (2015).

115. Parks, D. H. et al. Recovery of nearly 8,000 metagenome-assembled genomes substantially expands the tree of life. Nature Microbiology 2, 1533–1542 (2017).

116. Nurk S Meleshko D Korobeynikov A, P. P. metaSPAdes: A New Versatile Metagenomic Assembler. Genome Res. 27, 824–834 (2017).

117. Eren, A. M. et al. Anvi’o: An advanced analysis and visualization platformfor ’omics data. PeerJ 3, e1319 (2015).

118. Olm, M. R., Brown, C. T., Brooks, B. & Banfield, J. F. dRep: a tool for fast and accurate genomic comparisons that enables improved genome recovery from metagenomes through de-replication. ISME J. 11, 2864–2868 (2017).

119. Chaumeil, P.-A., Mussig, A. J., Hugenholtz, P. & Parks, D. H. GTDB-Tk: a toolkit to classify genomes with the Genome Taxonomy Database. Bioinformatics 36, 1925–1927 (2019).

120. Parks, D. H. et al. A standardized bacterial taxonomy based on genome phylogeny substantially revises the tree of life. Nat. Biotechnol. 36, 996–1004 (2018).

121. Hyatt, D. et al. Prodigal: Prokaryotic gene recognition and translation initiation site identification. BMC Bioinformatics 11, 119 (2010).

122. Eddy, S. R. Accelerated profile HMM searches. PLoS Comput. Biol. 7, e1002195 (2011).

123. Mistry, J. et al. Pfam: The protein families database in 2021. Nucleic Acids Res. 49, D412–D419 (2021).

124. Kanehisa, M. & Goto, S. KEGG: Kyoto Encyclopaedia of Genes and Genomes. Nucleic Acids Res. 28, 27–30 (2000).

125. Saheb Kashaf, S., et al. Integrating cultivation and metagenomics for a multi-kingdom view of skin microbiome diversity and functions. Nat Microbiol 7, 169–179 (2022).

126. Minich, J. J., et al. Quantifying and Understanding Well-to-Well Contamination in Microbiome Research. mSystems 4, (2019).

127. Murray, D. C., Coghlan, M. L. & Bunce, M. From benchtop to desktop: important considerations when designing amplicon sequencing workflows. PLoS One 10, e0124671 (2015).

128. Yu, Y., Lee, C., Kim, J. & Hwang, S. Group-specific primer and probe sets to detect methanogenic communities using quantitative real-time polymerase chain reaction. Biotechnol. Bioeng. 89, 670–679 (2005).

129. Binladen, J. et al. The use of coded PCR primers enables high-throughput sequencing of multiple homolog amplification products by 454 parallel sequencing. PLoS One 2, e197 (2007).

130. Schnell, I. B., Bohmann, K. & Gilbert, M. T. P. Tag jumps illuminated--reducing sequence-to-sample misidentifications in metabarcoding studies. Mol. Ecol. Resour. 15, 1289–1303 (2015).

131. Faircloth, B. C. & Glenn, T. C. Not all sequence tags are created equal: designing and validating sequence identification tags robust to indels. PLoS One 7, e42543 (2012).

132. Carøe, C. & Bohmann, K. Tagsteady: A metabarcoding library preparation protocol to avoid false assignment of sequences to samples. Mol. Ecol. Resour. 20, 1620–1631 (2020).

133. Martin, M. Cutadapt removes adapter sequences from high-throughput sequencing reads. EMBnet.journal 17, 10–12 (2011).

134. Callahan, B. J. et al. DADA2: High-resolution sample inference from Illumina amplicon data. Nat. Methods 13, 581–583 (2016).

135. Davis, N. M., Proctor, D. M., Holmes, S. P., Relman, D. A. & Callahan, B. J. Simple statistical identification and removal of contaminant sequences in marker-gene and metagenomics data. Microbiome 6, 226 (2018).

136. Ma, N. L. et al. Body mass, mercury exposure, biochemistry and untargeted metabolomics of incubating common eiders (*Somateria mollissima*) in three Baltic colonies. Environ. Int. 142, 105866 (2020).

137. Adusumilli, R. & Mallick, P. Data Conversion with ProteoWizard msConvert. in Proteomics: Methods and Protocols (eds. Comai, L., Katz, J. E. & Mallick, P.) 339–368 (Springer New York, 2017).

138. Pluskal, T., Castillo, S., Villar-Briones, A. & Oresic, M. MZmine 2: modular framework for processing, visualizing, and analyzing mass spectrometry-based molecular profile data. BMC Bioinformatics 11, 395 (2010).

139. Olivon, F., Grelier, G., Roussi, F., Litaudon, M. & Touboul, D. MZmine 2 Data-Preprocessing To Enhance Molecular Networking Reliability. Anal. Chem. 89, 7836– 7840 (2017).

140. Watrous, J. et al. Mass spectral molecular networking of living microbial colonies. Proc. Natl. Acad. Sci. U. S. A. 109, E1743–52 (2012).

141. Wang, M. et al. Sharing and community curation of mass spectrometry data with Global Natural Products Social Molecular Networking. Nat. Biotechnol. 34, 828–837 (2016).

142. Dührkop, K., Shen, H., Meusel, M., Rousu, J. & Böcker, S. Searching molecular structure databases with tandem mass spectra using CSI:FingerID. Proc. Natl. Acad. Sci. U. S. A. 112, 12580–12585 (2015).

143. Dührkop, K. et al. SIRIUS 4: a rapid tool for turning tandem mass spectra into metabolite structure information. Nat. Methods 16, 299–302 (2019).

144. Rasmussen, J. A. et al. A multi-omics approach unravels metagenomic and metabolic alterations of a probiotic and synbiotic additive in rainbow trout (*Oncorhynchus mykiss*). Microbiome 10, 21 (2022).

145. van der Hooft, J. J. J., Wandy, J., Barrett, M. P., Burgess, K. E. V. & Rogers, S. Topic modeling for untargeted substructure exploration in metabolomics. Proc. Natl. Acad. Sci. U. S. A. 113, 13738–13743 (2016).

146. Rogers, S. et al. Deciphering complex metabolite mixtures by unsupervised and supervised substructure discovery and semi-string-nameomated annotation from MS/MS spectra. Faraday Discuss. 218, 284–302 (2019).

147. da Silva, R. R. et al. Propagating annotations of molecular networks using *in silico* fragmentation. PLoS Comput. Biol. 14, e1006089 (2018).

148. Mohimani, H. et al. Dereplication of peptidic natural products through database search of mass spectra. Nat. Chem. Biol. 13, 30–37 (2017).

149. Djoumbou Feunang, Y., et al. ClassyFire: string-nameomated chemical classification with a comprehensive, computable taxonomy. J. Cheminform. 8, 61 (2016).

150. Ernst, M. et al. MolNetEnhancer: Enhanced Molecular Networks by Integrating Metabolome Mining and Annotation Tools. Metabolites 9, (2019).

151. Mock, A. et al. MetaboDiff: an R package for differential metabolomic analysis. Bioinformatics 34, 3417–3418 (2018).

152. Viant, M. R. et al. Use cases, best practice and reporting standards for metabolomics in regulatory toxicology. Nat. Commun. 10, 3041 (2019).

153. Ritchie, M. E. et al. limma powers differential expression analyses for RNA-sequencing and microarray studies. Nucleic Acids Res. 43, e47 (2015).

154. Lie, O. & Lambertsen, G. Fatty acid composition of glycerophospholipids in seven tissues of cod (*Gadus morhua*), determined by combined high-performance liquid chromatography and gas chromatography. J. Chromatogr. 565, 119–129 (1991).

155. Tastesen, H. S., Keenan, A. H., Madsen, L., Kristiansen, K. & Liaset, B. Scallop protein with endogenous high taurine and glycine content prevents high-fat, high-sucrose-induced obesity and improves plasma lipid profile in male C57BL/6J mice. Amino Acids 46, 1659–1671 (2014).

156. Mallick, H., et al. Multivariable Association Discovery in Population-scale Meta-omics Studies. bioRxiv 2021.01.20.427420 (2021) doi:10.1101/2021.01.20.427420.

157. Alberdi, A. & Gilbert, M. T. P. A guide to the application of Hill numbers to DNA-based diversity analyses. Mol. Ecol. Resour. 19, 804–817 (2019).

158. McMurdie, P. J. & Holmes, S. phyloseq: an R package for reproducible interactive analysis and graphics of microbiome census data. PLoS One 8, e61217 (2013).

159. Zhou, X. & Stephens, M. Genome-wide efficient mixed-model analysis for association studies. Nat. Genet. 44, 821–824 (2012).

160. Meisner, J. & Albrechtsen, A. Inferring population structure and admixture proportions in low-depth NGS data. Genetics 210, 719–731 (2018).

161. Quinlan, A. R. & Hall, I. M. BEDTools: A flexible suite of utilities for comparing genomic features. Bioinformatics 26, 841–842 (2010).

162. Ahlmann-Eltze, C. & Patil, I. ggsignif: R Package for Displaying Significance Brackets for ‘ggplot2’. (2021) doi:10.31234/osf.io/7awm6.

163. Buchfink, B., Xie, C. & Huson, D. H. Fast and sensitive protein alignment using DIAMOND. Nat. Methods (2015) doi:10.1038/nmeth.3176.

164. Benedict, M. N., Henriksen, J. R., Metcalf, W. W., Whitaker, R. J. & Price, N. D. ITEP: an integrated toolkit for exploration of microbial pan-genomes. BMC Genomics 15, 8 (2014).

165. van Dongen, S. & Abreu-Goodger, C. Using MCL to Extract Clusters from Networks. in Bacterial Molecular Networks: Methods and Protocols (eds. van Helden, J., Toussaint, A. & Thieffry, D.) 281–295 (Springer New York, 2012).

166. Price, M. N., Dehal, P. S. & Arkin, A. P. FastTree: computing large minimum evolution trees with profiles instead of a distance matrix. Mol. Biol. Evol. 26, 1641–1650 (2009).

167. Pritchard, L., Glover, R. H., Humphris, S., Elphinstone, J. G. & Toth, I. K. Genomics and taxonomy in diagnostics for food security: soft-rotting enterobacterial plant pathogens. Anal. Methods 8, 12–24 (2015).

168. Barbosa, M. S. et al. Host cell interactions of novel antigenic membrane proteins of *Mycoplasma agalactiae*. BMC Microbiol. 22, 93 (2022).

169. Harkey, C. W., Everiss, K. D. & Peterson, K. M. The *Vibrio cholerae* toxin-coregulated-pilus gene tcpI encodes a homolog of methyl-accepting chemotaxis proteins. Infect. Immun. 62, 2669–2678 (1994).

170. Rivas, A. J., Lemos, M. L. & Osorio, C. R. *Photobacterium damselae* subsp. *damselae*, a bacterium pathogenic for marine animals and humans. Front. Microbiol. 4, 283 (2013).

171. Madden, T. L. et al. BLAST+: architecture and applications. BMC Bioinformatics 10, 421 (2009).

