## Supplementary Figures for "Host-gut microbiota interactions shape parasite infections in farmed Atlantic salmon"

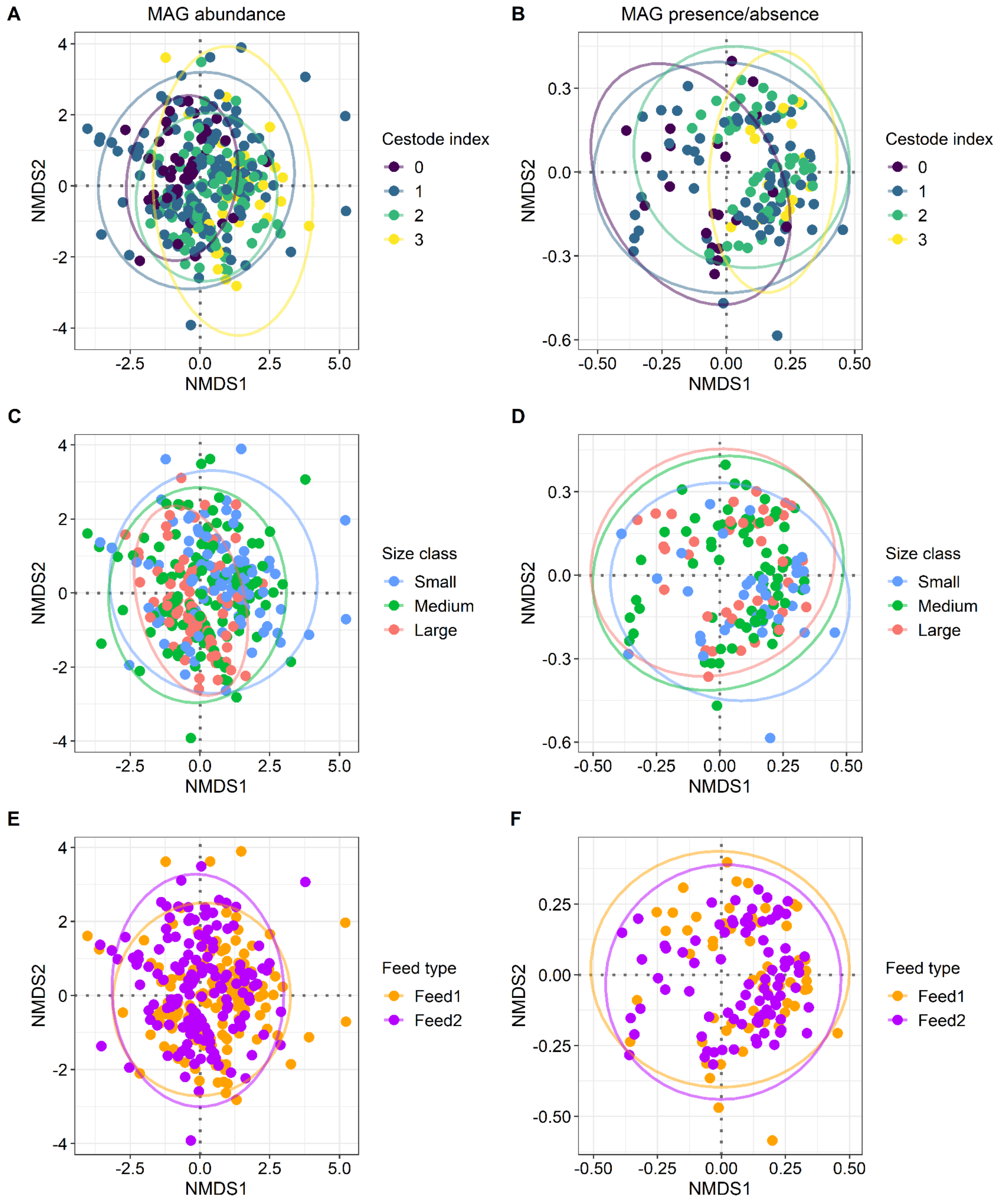


Supplementary Figure 1. Ordination (NMDS) of metagenomes, based on MAG abundance (A, C, E) and MAG presence/absence (B, D, F). Individuals are coloured by cestode index (A, B), size class (C, D) or feed type (E, F). For MAG abundance (A, C, E), the NMDS was generated using Euclidean distances (NMDS stress: 0.200, k = 2). Based on PERMANOVA results, cestode index contributed to 5.70% of the variation in abundance (F = 8.21, p = 0.001), size class to 3.56% (F = 7.70, p = 0.001), and feed type to 1.68% (F = 7.28, p = 0.001). For presence/absence (B, D, F), the NMDS was generated from Jaccard distances

(NMDS stress: 0.135, k = 3). In the PERMANOVA, cestode index contributed to 7.53% of the presence/absence of MAGs ($F = 10.87$, $p = 0.001$), size class to 2.11% ($F = 4.57$, $p = 0.001$) and feed type to 1.50% ($F = 6.48$, $p = 0.001$). Detailed results from the PERMANOVAs are presented in Supplementary Table 3.

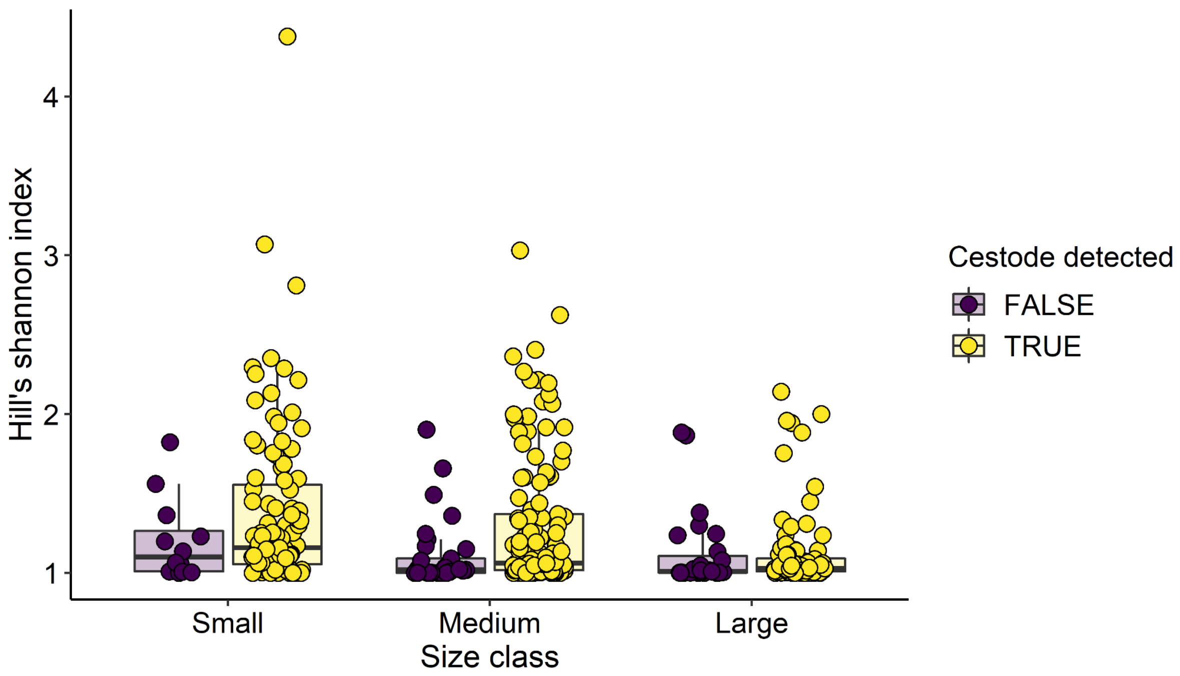


Supplementary Figure 2. Alpha diversity (Hill's Shannon index) of metagenomes, based on MAG abundance, stratified by cestode detection and size class.

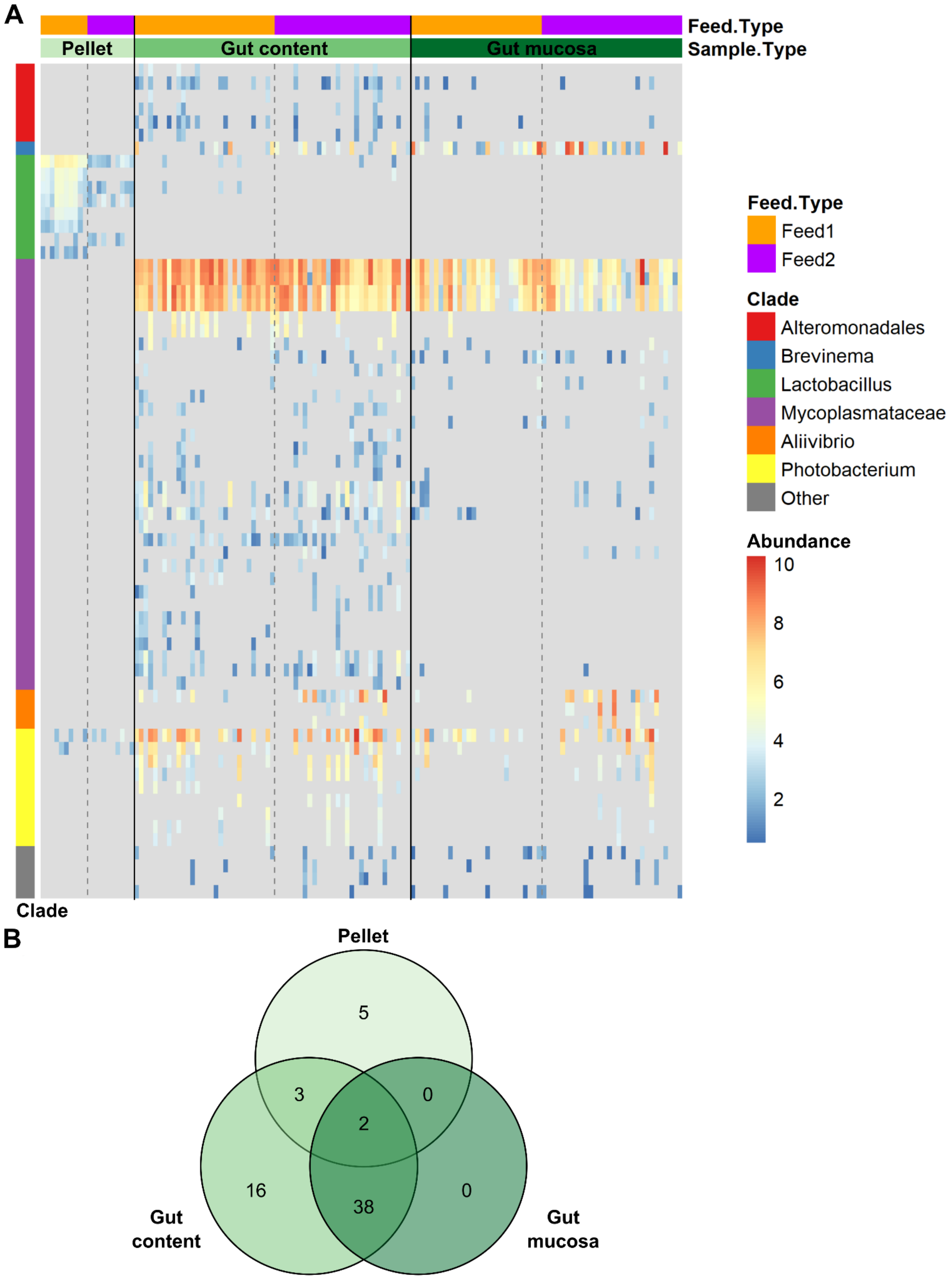


Supplementary Figure 3. Microbiota community composition of salmon gut content (same sample type as used for MAG analysis) compared to salmon gut mucosa scrapes and feed pellet, determined by 16S metabarcoding. **A)** Heatmap of the abundance (centre-log-ratio normalised) of the 64 amplicon sequence variants (ASVs) present in at least 5 samples after quality control. ASVs are annotated by their taxonomic clade, where “other” indicates ASVs that could not be assigned taxonomy at the level of order. ASVs within the order

*Alteromonadales* were classified as the genus *Neiella*. ASVs within the family *Mycoplasmataceae* were classified as the genus *Mycoplasma* (17 ASVs) or an unknown genus within the *Mycoplasmataceae* family (13 ASVs). Other taxonomic clades are provided at the genus level. Samples are annotated by sample type and commercial feed type. **B)** Venn diagram of the 64 ASVs in A detected at least once within each sample type. The 16 ASVs unique to gut content samples included 14 low-abundant *Mycoplasmataceae* taxa and 2 *Alteromonadales* taxa.

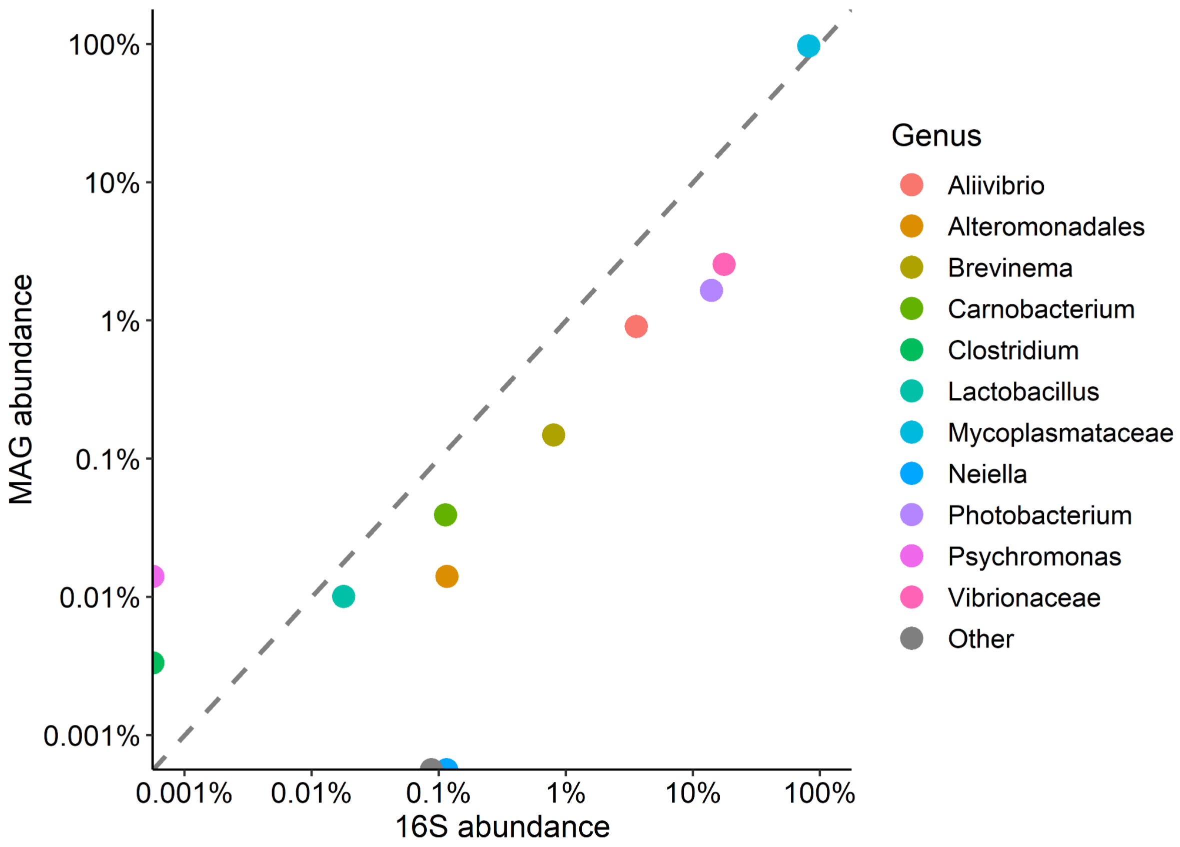


Supplementary Figure 4. Comparison of microbial relative abundances in salmon gut content samples captured by MAG assembly versus 16S profiling. The colours represent microbial genera, families, or orders in the case of Alteromonadales, as defined in the legend. The dashed line indicates hypothetical perfect concordance in relative abundances between the MAG and 16S profiling methods. Relative abundances are shown on a log10 scale. Generally, good agreement was observed for the most common microbial taxa. However, *Psychromonas* and *Clostridium* were only detected by MAG assembly (with relative abundances < 0.1%), while *Neiella* was only detected by 16S profiling. *Psychromonas* and *Neiella* are both of the order Alteromonadales and have somewhat similar relative abundances in the MAG and 16S datasets, respectively, suggesting that they may partly represent the same cluster of microbes that were assigned different taxonomies at lower levels due to differences in the databases used. ‘Other’ represents all 16S amplicon sequencing variants (ASVs) that are not represented by the MAG catalogue and accounts for approximately 0.1% of the total number of reads assigned to ASVs.

### Host variables

Cestode detection

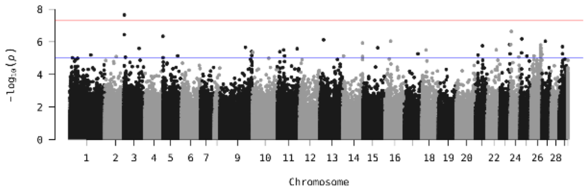


Gutted weight

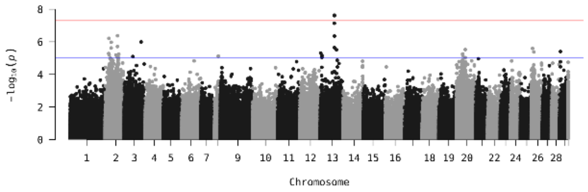


Feed type

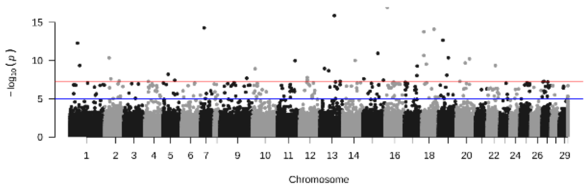


### MAG detection

MAG02 *Mycoplasma* sp.

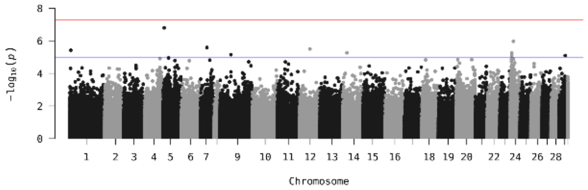


MAG03 *Mycoplasma* sp.

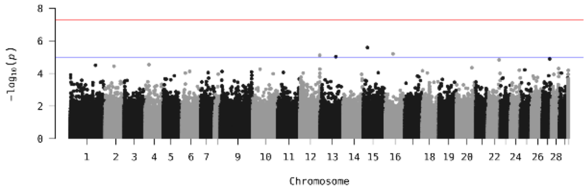


MAG04 *Mycoplasma* sp.

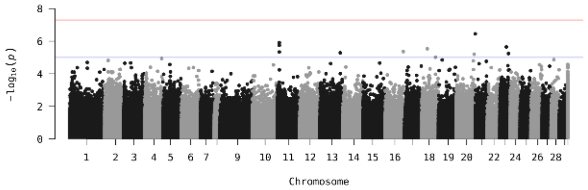


MAG05 *Mycoplasma* sp.

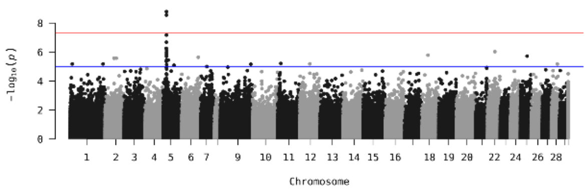


MAG06 *P. phosphoreum*

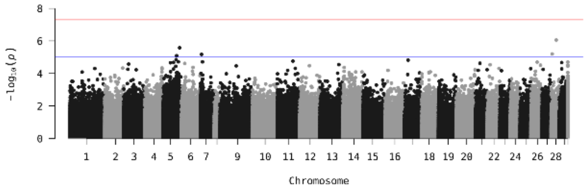


MAG07 *P. iliopiscarium*

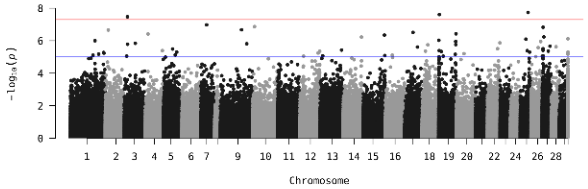


MAG08 *Aliivibrio* sp.

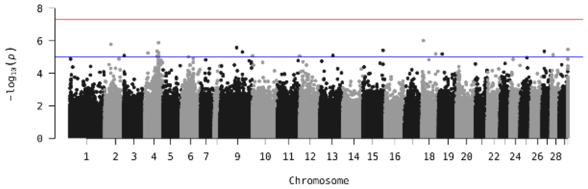


MAG09 *Aliivibrio* sp.

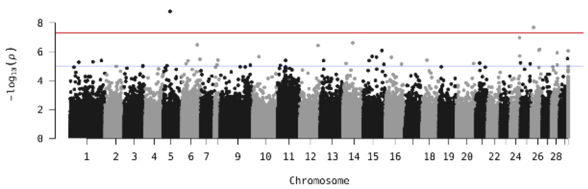


MAG11 *Brevinema* sp.

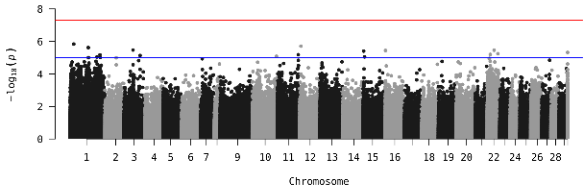


MAG12 *C. maltaromaticum*

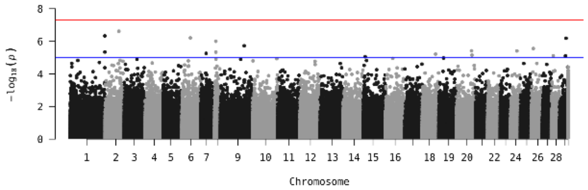


MAG13 *L. johnsonii*

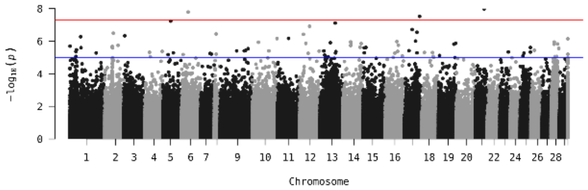

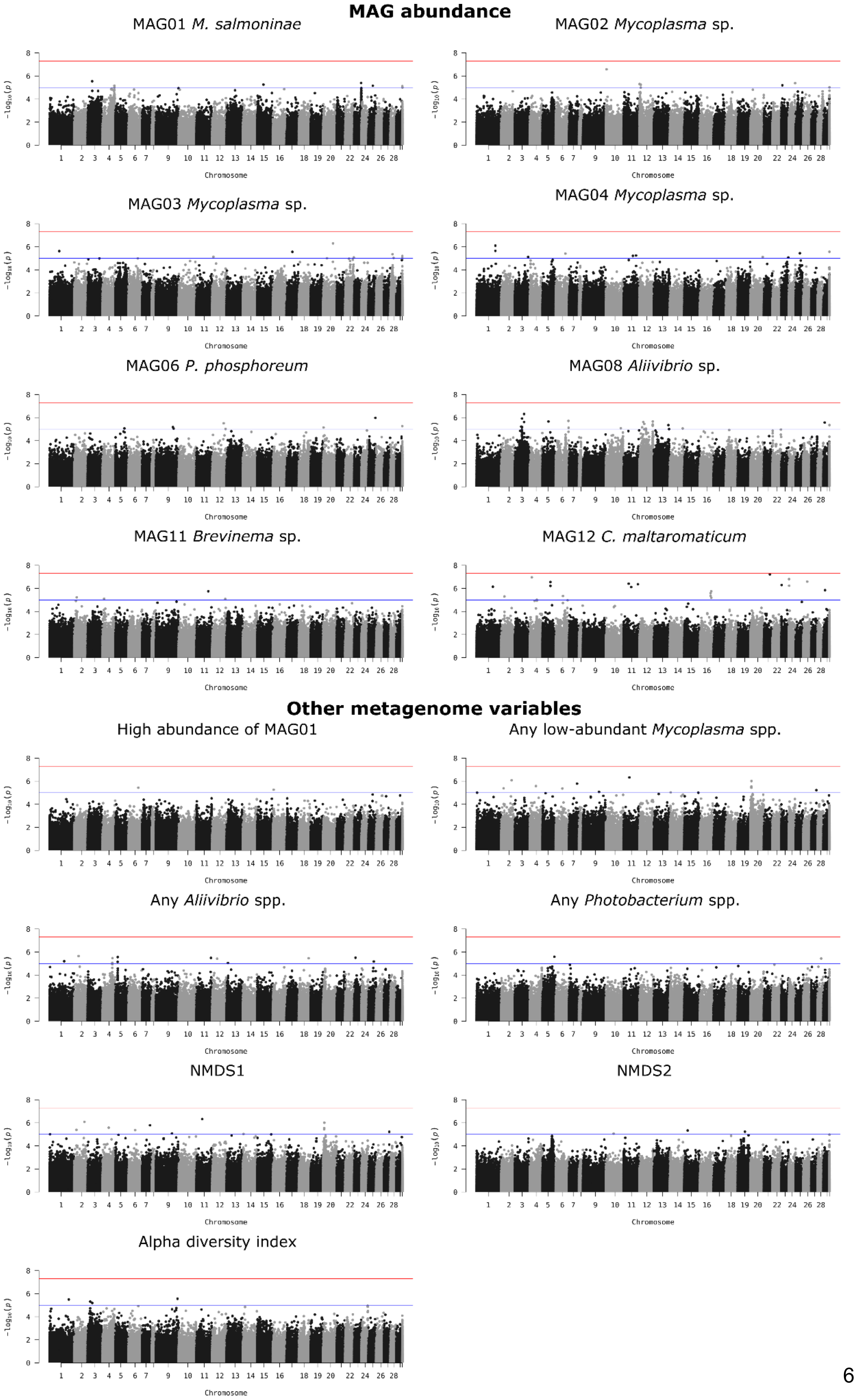
MAG abundance
MAG01 M. salmoninae
MAG02 Mycoplasma sp.
MAG03 Mycoplasma sp.
MAG04 Mycoplasma sp.
MAG06 P. phosphoreum
MAG08 Aliivibrio sp.
MAG11 Brevinema sp.
MAG12 C. maltaromaticum
Other metagenome variables
High abundance of MAG01
Any low-abundant Mycoplasma spp.
Any Aliivibrio spp.
Any Photobacterium spp.
NMDS1
NMDS2
Alpha diversity index
Chromosome
−log10(p)

Supplementary Figure 5. Manhattan plots of all GWAS runs. P-values of individual SNPs have been transformed on a -log10 axis. Red lines indicate the threshold for genome-wide significance ($p < 5e-8$) and blue lines indicate the suggestive threshold ($p < 1e-5$). Results of GWAS are summarised in Supplementary Table 6.

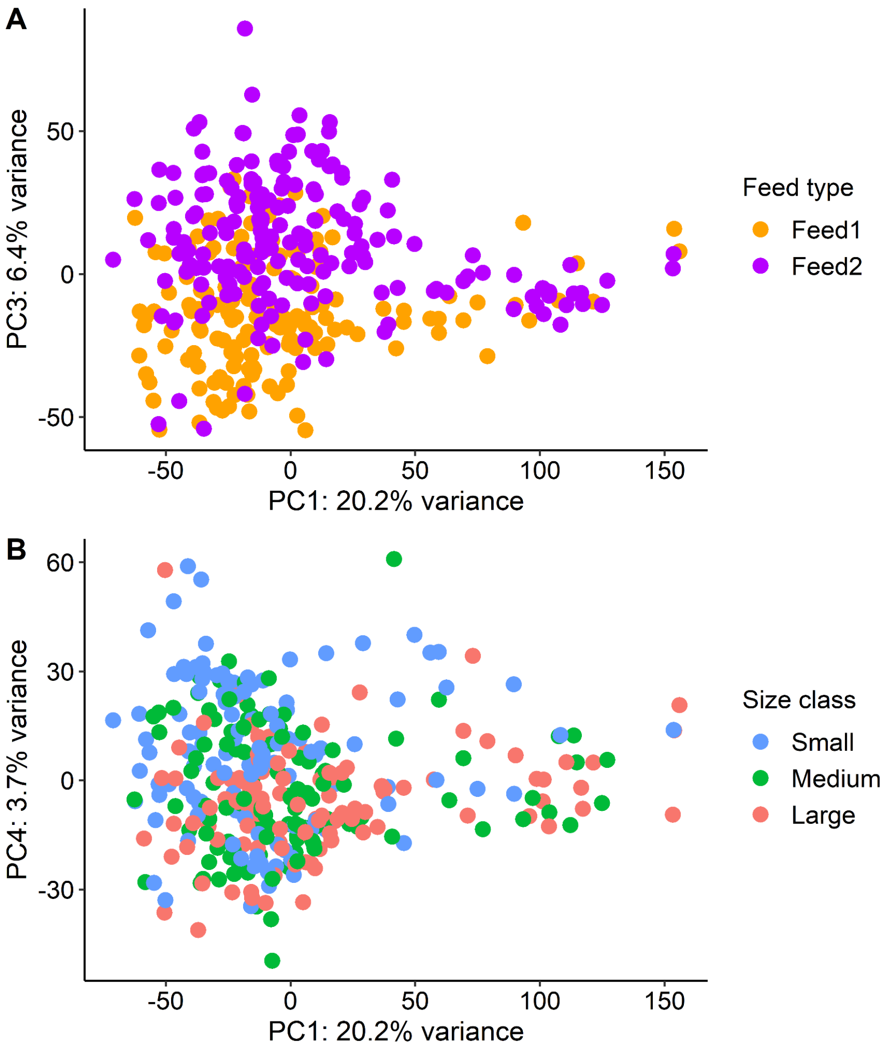


Supplementary Figure 6. Ordination (PCA) of host gut epithelial transcriptomes. Individuals showed some variation by feed type on PC3 (**A**) and some variation by size class on PC4 (**B**). PCAs were based on normalised and transformed gene expression data. Based on PERMANOVA results, feed type contributed to 2.7% of the variation in gene expression values ($F = 9.63$, $p = 0.001$) and size class contributed to 2.0% of the variation ($F = 3.53$, $p = 0.001$). Detailed results from the PERMANOVAs are presented in Supplementary Table 3.

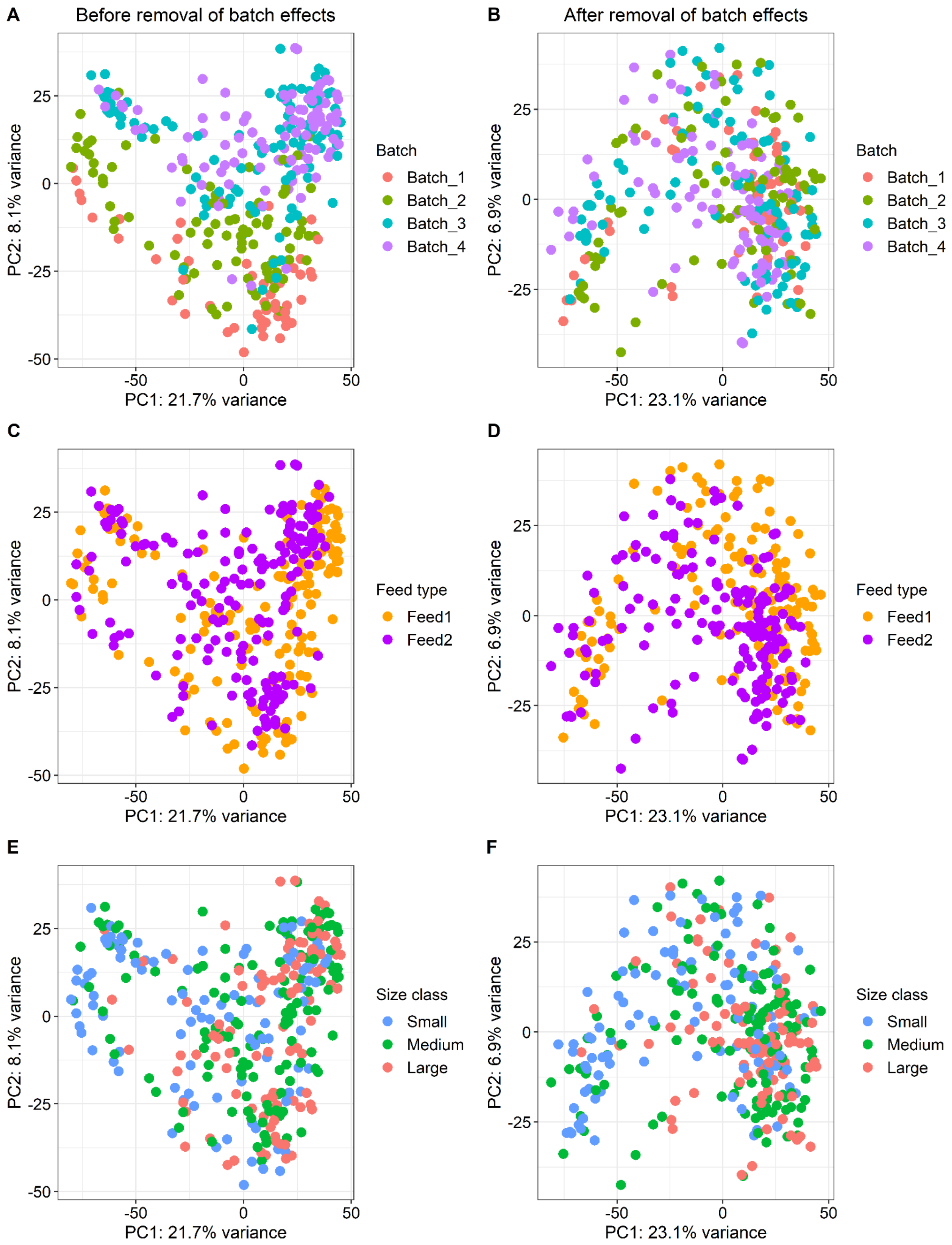


Supplementary Figure 7. Ordination (PCA) of gut metabolomes, before (**A, C, E**) and after (**B, D, F**) removal of batch effects. Batch effects are a common problem in metabolomic studies and in this study, metabolome batch explains 11.4% of the variation in metabolite abundance before correction ($F = 14.88$, $p = 0.001$) (**A**), while after correction, batch does not significantly explain metabolite abundance variation (**B**). Feed type consistently explains 2.2-2.5% of the variation ($F = 8.56$, $p = 0.001$) (**C–D**) and size class explains 3.2-3.6% of the variation ($F = 6.35$, $p = 0.001$). Detailed results from the PERMANOVAs are presented in Supplementary Table 3.

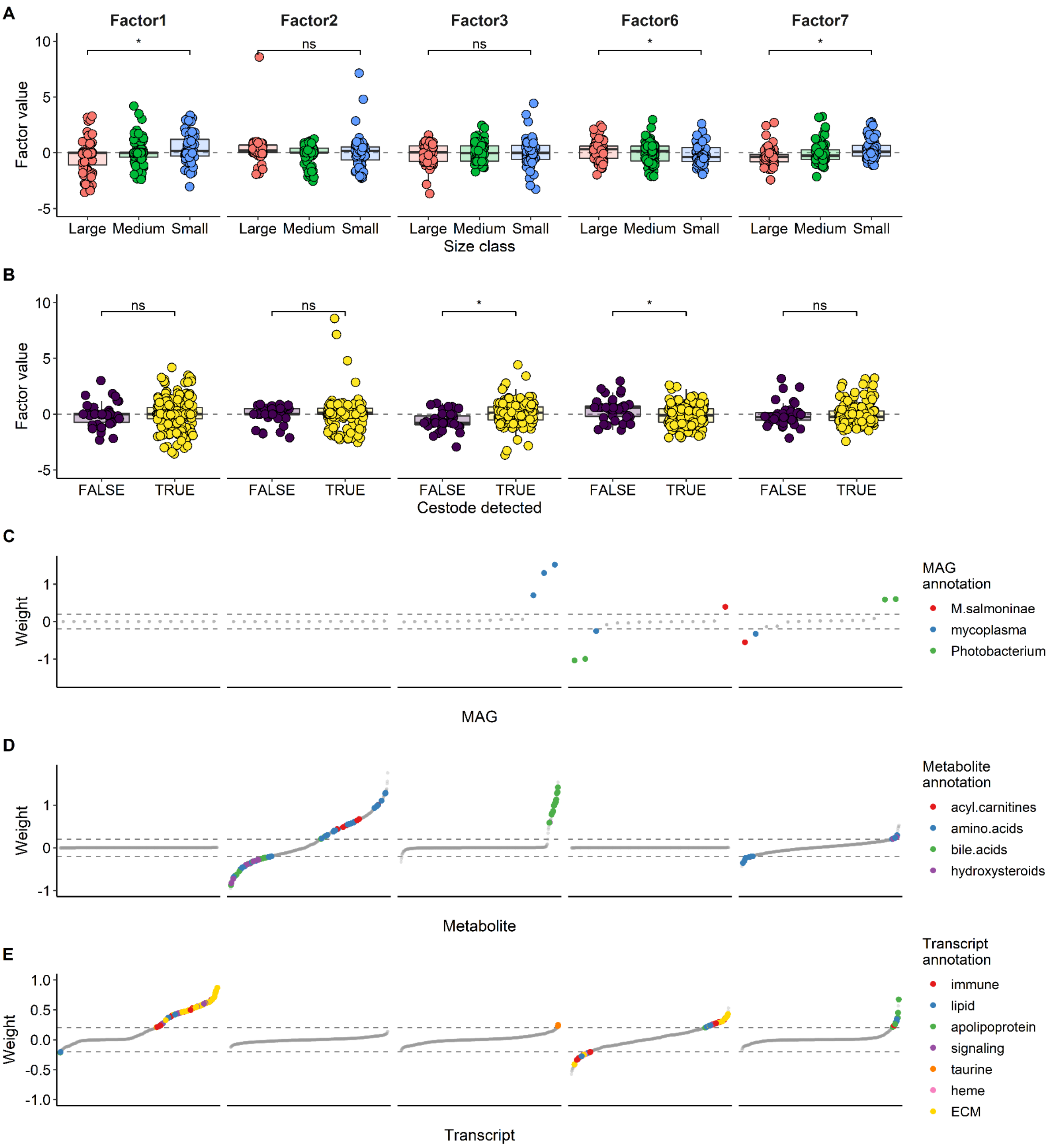


Supplementary Figure 8. MOFA results from the **Feed1** model, for Factors 1, 2, 3, 6 and 7. These factors from the Feed1 model are shown here because they captured similar patterns in variation as in the full combined model (Figure 4). The factor numbers are not comparable between models (for links between factors in each model, see Supplementary Table 8). **A**) Factors 1, 6 and 7 were correlated with size class. **B**) Factors 3 and 6 were correlated with cestode detection. **C–E**) Feature weights for the metagenome (**C**), metabolome (**D**) and transcriptome (**E**) for Factors 1, 2, 3, 6 and 7. Features are ranked according to their weight. The higher the absolute weight, the more strongly associated a feature is with that factor. A positive weight indicates the feature has higher levels in samples with positive factor values, while a negative weight indicates the opposite. Features with weights > 0.2 or < -0.2 are coloured by MAG species or genus (C) or most frequent functional annotation (D-E), while features with less frequent annotations, or those with weights outside the threshold range (as indicated by the dashed lines) are shown in grey.

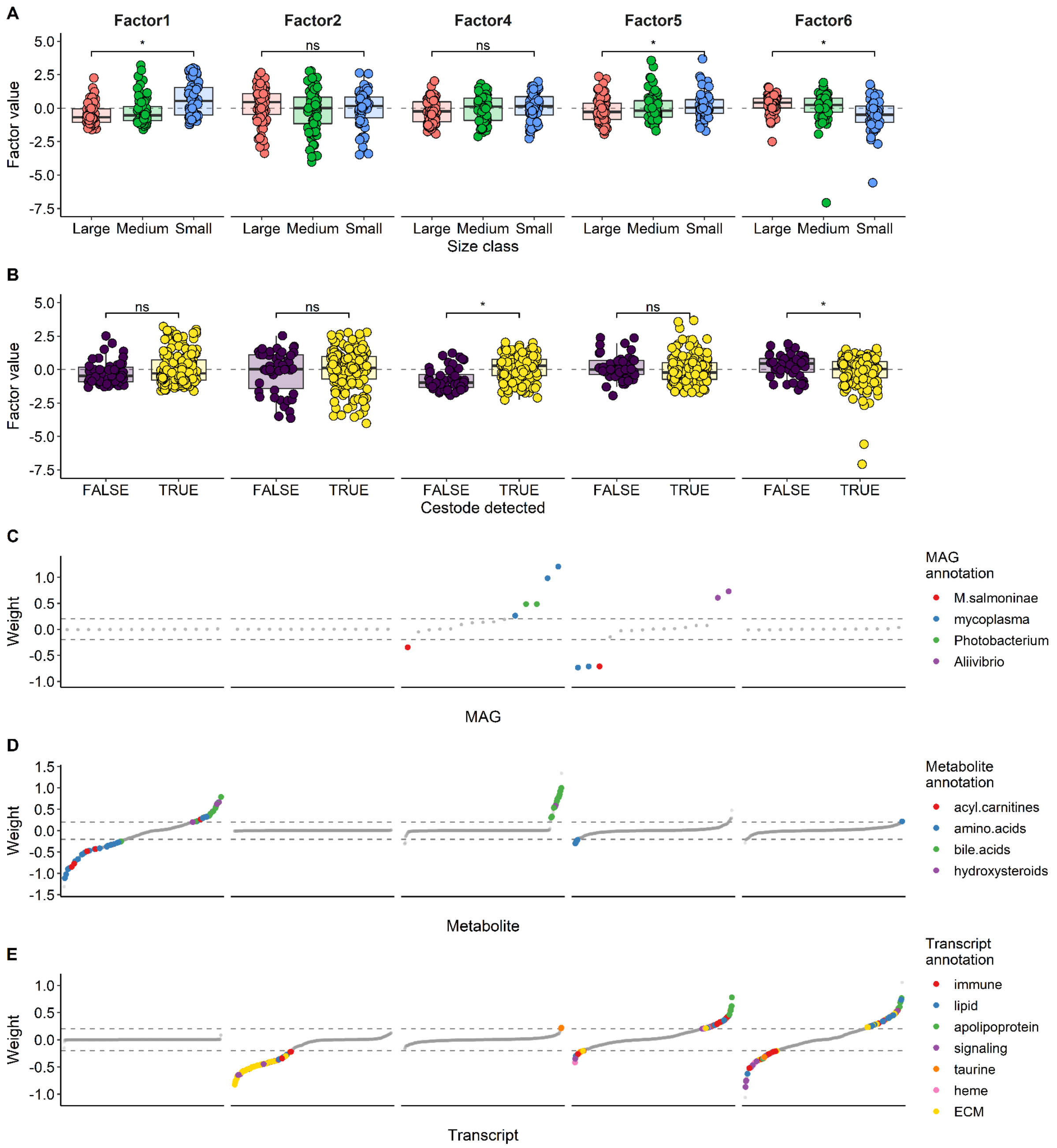


Supplementary Figure 9. MOFA results from the **Feed2** model, for Factors 1, 2, 4–6. These factors from the Feed2 model are shown here because they captured similar patterns in variation as in the full combined model (Figure 4). The factor numbers are not comparable between models (for links between factors in each model, see Supplementary Table 8). **A**) Factors 1, 5 and 6 were correlated with size class. **B**) Factors 4 and 6 were correlated with cestode detection. **C–E**) Feature weights for the metagenome (**C**), metabolome (**D**) and transcriptome (**E**) for Factors 1, 2, 4–6. Features are ranked according to their weight. The higher the absolute weight, the more strongly associated a feature is with that factor. A positive weight indicates the feature has higher levels in samples with positive factor values, while a negative weight indicates the opposite. Features with weights > 0.2 or < -0.2 are coloured by MAG species or genus (C) or most frequent functional annotation (D-E), while features with less frequent annotations, or those with weights outside the threshold range (as indicated by the dashed lines) are shown in grey.
